# Robust and Functional Immunity up to 9 months after SARS-CoV-2 infection: a Southeast Asian longitudinal cohort

**DOI:** 10.1101/2021.08.12.455901

**Authors:** Vo Hoa Thi My, Maestri Alvino, Auerswald Heidi, Sorn Sopheak, Lay Sokchea, Heng Seng, Sann Sotheary, Ya Nisa, Pean Polidy, Dussart Philippe, Schwartz Olivier, Ly Sovann, Bruel Timothee, Ly Sowath, Duong Veasna, Karlsson Erik A, Cantaert Tineke

**Affiliations:** Immunology Unit, Institut Pasteur du Cambodge, Institut Pasteur International Network, Phnom Penh, Cambodia; Virology Unit, Institut Pasteur du Cambodge, Institut Pasteur International Network, Phnom Penh, Cambodia; Epidemiology and Public Health Unit, Institut Pasteur du Cambodge, Institut Pasteur International Network, Phnom Penh, Cambodia; Cambodian CDC-MoH; Virus & Immunity Unit, Department of Virology, Institut Pasteur, CNRS UMR3569, Paris, France

**Author notes:** co-first authors (Vo Hoa Thi My, Maestri Alvino). co-last authors (Karlsson Erik A, Cantaert Tineke).

## Abstract

Assessing the duration of humoral and cellular immunity remains key to overcome the current SARS-CoV-2 pandemic, especially in understudied populations in least developed countries. Sixty-four Cambodian individuals with laboratory-confirmed infection with asymptomatic or mild/moderate clinical presentation were evaluated for humoral immune response to the viral spike protein and antibody effector functions during acute phase of infection and at 6-9 months follow-up. Antigen-specific B cells, CD4^+^ and CD8^+^ T cells were characterized, and T cells were interrogated for functionality at late convalescence. Anti-spike (S) antibody titers decreased over time, but effector functions mediated by S-specific antibodies remained stable. S- and nucleocapsid (N)-specific B cells could be detected in late convalescence in the activated memory B cell compartment and are mostly IgG^+^. CD4^+^ and CD8^+^ T cell immunity was maintained to S and membrane (M) protein. Asymptomatic infection resulted in decreased ADCC and frequency of SARS-CoV-2-specific CD4^+^ T cells at late convalescence. Whereas anti-S antibodies correlated with S-specific B cells, there was no correlation between T cell response and humoral immunity. Hence, all aspects of a protective immune response are maintained up to nine months after SARS-CoV-2 infection in the absence of re-infection.

**One sentence summary:** Functional immune memory to SARS-CoV-2, consisting of polyfunctional antibodies, memory B cells and memory T cells are maintained up to nine months in a South-East Asian cohort in the absence of re-infection.

## Introduction

In December 2019, a cluster of severe pneumonia of unknown cause was reported to the World Health Organization. Investigation into the etiology revealed a novel betacoronavirus, subsequently named Severe Acute Respiratory Syndrome Coronavirus 2 (SARS-CoV-2), the causative agent of Coronavirus Disease 2019 (COVID-19) (1). Clinical spectrum of COVID-19 ranges from asymptomatic, over mild upper respiratory tract illness, to severe viral pneumonia resulting in respiratory failure and death (1–3).

Upon infection with SARS-CoV-2, humans generate SARS-CoV-2-specific antibodies, memory B cells, and CD4^+^ and CD8^+^ T cells, which all have complementary functions in the clearance of SARS-CoV-2 virions and infected cells (4). Mainly structural proteins are targeted by the immune response, such as the membrane (M) and spike (S) protein integrated in the virion envelope, and the nucleoprotein (N), which protects the RNA genome (5–7). The S protein consists of two domains. The S1 region contains the receptor binding domain (RBD) which interacts with the host protein Angiotensin-converting enzyme 2 (ACE2) to mediate cell entry, whereas the S2 domain mediates membrane fusion. The S1 domain with the RBD is a major target of neutralizing antibodies (8, 9). Several studies show correlation between antibodies targeting S and functional neutralization (10–12). In animal models, these neutralizing antibodies are protective against secondary infection (13, 14). In humans, anti-S antibodies and neutralizing antibodies can be detected up to one year post infection (15–17).

Besides neutralization, antibodies activate a variety of effector functions mediated by their Fc domain. These include complement activation, killing of infected cells and phagocytosis of viral particles (18). Indeed, it has been shown that symptomatic and asymptomatic SARS-CoV-2 infection elicit polyfunctional antibodies targeting infected cells (19, 20) and Fc mediated effector activity of antibodies correlates with reduced disease severity and mortality after SARS-CoV-2 infection (21). However, the evolution of this response over time requires further investigation (22, 23).

Persistence of serum antibodies may not be the sole determinant of long-lasting immunity post infection or vaccination. Anamnestic recall of memory T and B cell populations can also reduce infection or disease at re-exposure (24–26), with increasing importance as antibody titers wane. Virus-specific memory T and B cells can be detected in at least 50% of the individuals six months post infection (24, 26, 27). Several studies suggest that increased severity of COVID-19 induces a stronger SARS-CoV-2-specific CD4^+^ T cell response (27–29). However, the magnitude, quality, and protective capacity of cellular responses against SARS-CoV-2 requires further definition (4).

Kinetics and duration of the memory immune responses could depend on a number of factors including viremia, disease severity, re-infection, cross-reactivity with human seasonal coronaviruses (hCoVs), ethnic background, and age (reviewed in (4)). Other human betacoronaviruses, such as hCoV OC43 and HKU1, and zoonotic viruses, such as SARS-CoV-1 and Middle East respiratory syndrome-related coronavirus (MERS-CoV), show waning antibody levels as soon as three months post infection. In contrast, T cell responses are detectable up to 17 years later (30, 31).

Most studies analyzing the evolution of the adaptive immune response to SARS-CoV-2 are conducted in Caucasian populations (4). In South-East Asia, very few studies have been performed, which mainly focused on antibody responses (16, 32–34). Understanding long-term immunity after natural infection by determining the frequency, function, and specificities of the humoral and cellular immune components in various populations is critical. Paucity of data from at risk areas and populations can hamper global mitigation and vaccination efforts.

We comprehensively characterized long-lived immunity in 64 Cambodian individuals with laboratory-confirmed infection experiencing mild/moderate or asymptomatic clinical outcome. Cambodia remained almost completely COVID-19-free in 2020 (35), hence, additional exposure to SARS-CoV-2 in this cohort is highly unlikely. The humoral immune response to the viral spike protein was assessed and antibody effector functions were characterized during the acute phase of infection and up to nine months later. In addition, at late convalescence, persistence and phenotype of S1- and N-specific memory B cells was evaluated. Virus-specific CD4^+^ and CD8^+^ T cells were characterized and T cells were interrogated for functionality.

## Results

### Long-term follow-up of SARS-CoV-2 imported cases

Sixty-four individuals with confirmed SARS-CoV-2 were included and re-assessed 6-9 months after infection. SARS-CoV-2 infection was confirmed by positive molecular diagnosis as part of the national surveillance system. Since Cambodia had minimal detection of SARS-CoV-2 during the follow-up period, the probability of re-exposure to SARS-CoV-2 was minimal (35) in 2020. For 33 individuals, we obtained a blood sample 2-9 days after laboratory confirmed infection (Figure S1A). For all 64 study participants, between 1 to 15 follow-up nasopharyngeal/oropharyngeal (NP/OP) swab samplings assessed the duration of viremia during the acute phase of infection via RT-PCR (36). Based on the duration of viremia, 53% of individuals were considered “long shedders” with detection of viral RNA in NP/OP swabs for ≥10 days (Figure S1B). Overall, 70% of the patients displayed mild or moderate symptoms, and 30% remained asymptomatic (Table S1). For all assays, samples were selected based on availability and quality.

### Asymptomatic and mild/moderate infection induces a persisting anti-spike antibody response

The presence of S-binding antibodies was measured using the S-Flow assay, which sensitively and quantitatively measures anti-S IgG, IgA, and IgM by flow cytometry (19, 37) (Figure 1A). The National Institute for Biological Standards and Control (NIBSC) references were utilized to validate the assays and pre-pandemic samples obtained from nineteen individuals were measured to set the cutoff for each assay (Figure S2). Anti-S IgM, IgG, and IgA titers decreased significantly between acute phase and late convalescence (p=0.02, p<0.0001, p<0.0001, respectively). (Figure 1B). Within the total S-binding antibodies, the percentage of anti-S IgM and IgA decreased whereas anti-S IgG increased over time (p=0.0003, Figure 1C). The detection of neutralizing antibodies was achieved by foci reduction neutralization test using a Cambodian SARS-CoV-2 isolate. There was no difference in the titers of SARS-CoV-2 neutralizing antibodies between the acute and convalescent phase, even though titers tended to decrease over time (Figure 1D). Over time, the percentage of individuals positive for anti-S IgM (p<0.0001) and anti-S IgA (p<0.0001) decreased (Figure 1E). In the acute phase, 91% of individuals were positive for anti-S IgG, and only 70% of the individuals were positive for neutralizing antibody titers. Up to nine months post infection, the frequency of individuals positive for anti-S IgG remained stable (88%) whereas the frequency of individuals with neutralizing titers decreased to 56% (p=0.055) (Figure 1E). Analyzing only individuals with paired samples available, revealed similar results as the whole cohort (Figure S3, A-B). Taken together, these data show that despite decreases in antibody titers over time, the percentage of individuals positive for anti-S IgG remains stable.

**Figure 1.**
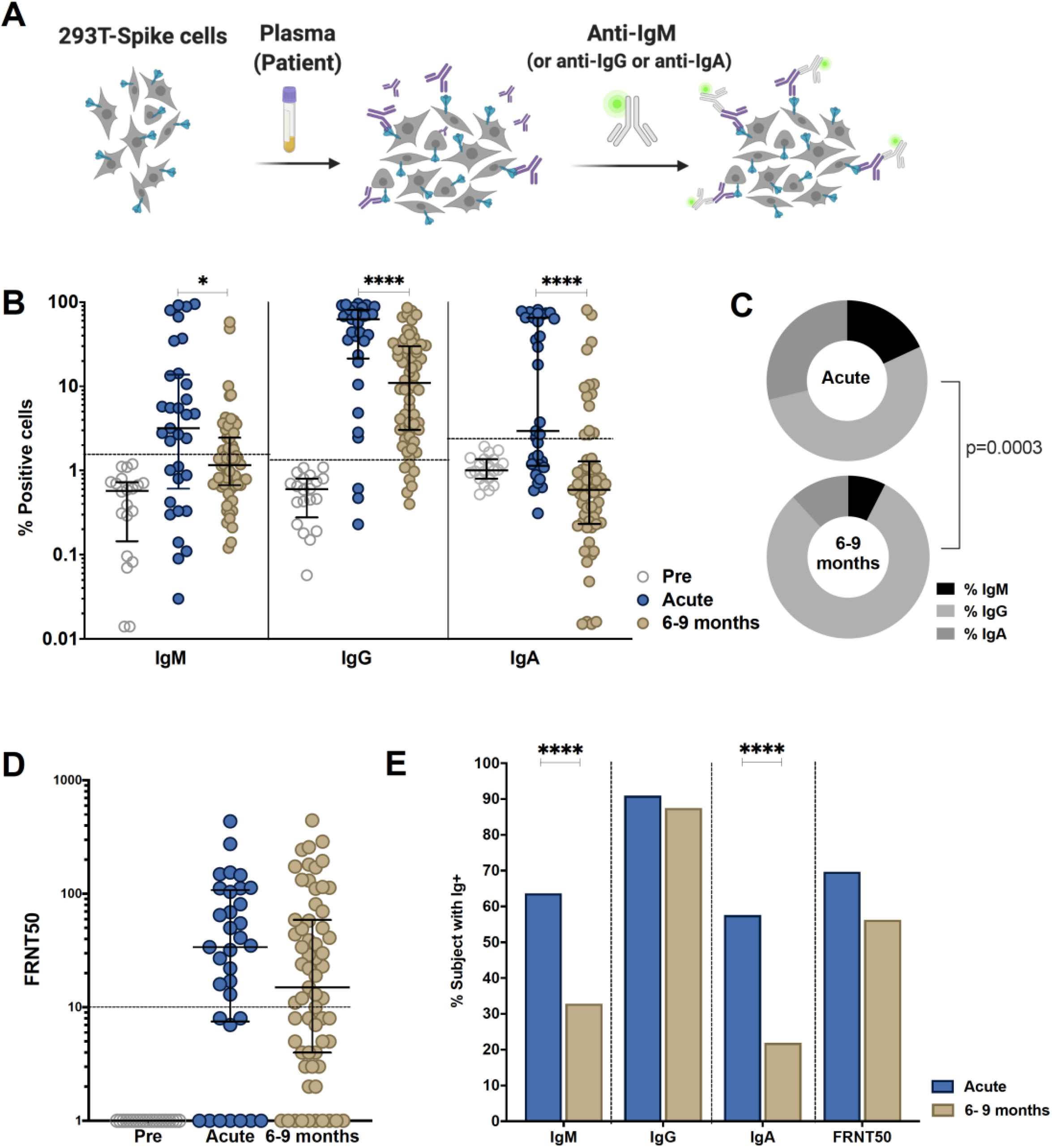
Comparison of antibody response in SARS-CoV-2-infected individuals during the acute phase and 6-9 months post infection. Individuals were sampled 2-9 days post laboratory confirmation and 6-9 months later. (**A**) Schematic model of the S-Flow assay. (**B**) Amount of antibodies against spike protein were reported as percentage of spike-expressing 293T cells bound by IgM, IgG, IgA in the S-Flow assay. (**C**) Pie charts show the proportion of anti-S IgM, IgG and IgA antibodies. (**D**) SARS-CoV-2 neutralizing activity was calculated as FRNT50 titer in foci reduction neutralization test (FRNT). (**E**) Comparison of the percentage of individuals positive for anti-S IgM, IgG, IgA and FRNT50. Statistical comparisons were performed by Mann Whitney test (B and D) and Chi-square test (C and E). The dashed line indicates the cutoff for positivity based on values calculated following formula: cut-off = % mean positive cells from 19 pre-pandemic samples + 3x standard deviation. Each dot represents the result from a single individual. Lines represent median and IQR. *p < 0.05, **p <0.01, ***p <0.001, and ****p < 0.0001. Pre-pandemic n=19, acute n=33, 6-9 months n=64.

### Functional antibody response changes over time post SARS-CoV-2 infection

Besides neutralization, antibodies can mediate Fc-effector functions, such as complement activation, killing of virus-infected cells and phagocytosis of viral particles (18). To further define the humoral response in these individuals, we assessed antibody effector functions *in vitro*. The NIBSC references were utilized to validate the assays and nineteen pre-pandemic samples were measured to set the cutoff for each assay. Antibody-dependent cellular phagocytosis (ADCP) assay measures the engulfment of neutravidin beads coated with SARS-CoV-2 derived S1 by THP-1 cells (Figure 2A, S4). A decrease in ADCP can be observed between the acute and late convalescent phase (p=0.005, Figure 2B, C). The percentage of subjects with ADCP activity decreased from 73% to 55% over time. However, when calculating the proportion of ADCP within the total anti-S antibodies, we observed a significant increase of the proportion of ADCP over time (p=0.003, Figure 2D).

**Figure 2.**
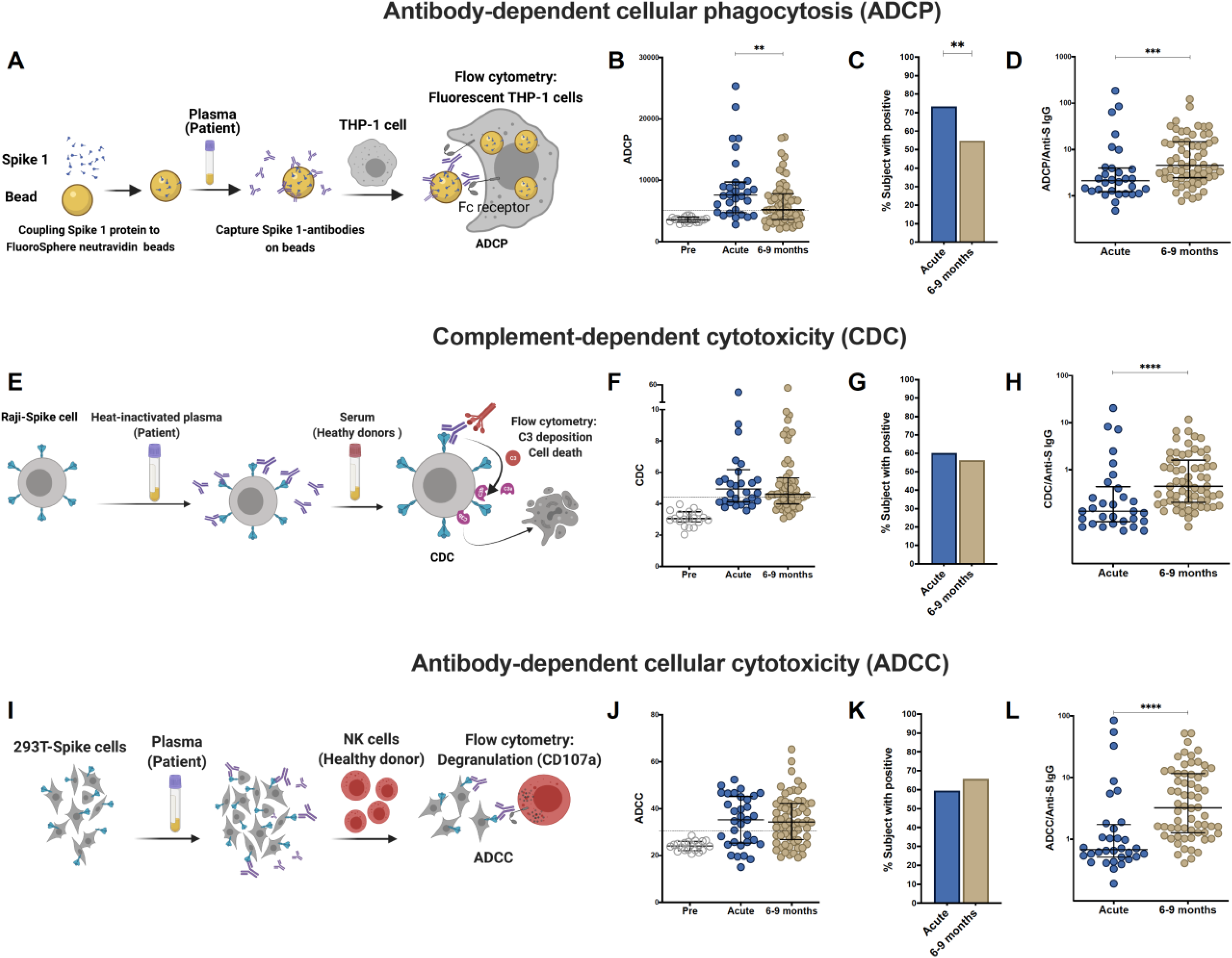
Comparison of effector function profiles of plasma from SARS-CoV-2-infected individuals during the acute phase and 6-9 months post infection. Individuals were sampled 2-9 days post laboratory confirmation and 6-9 months later. (**A**) Schematic representation of the antibody-dependent cellular phagocytosis (ADCP) assay. (**B**) Comparison of ADCP activity in pre-pandemic samples, SARS-CoV-2-infected individuals in the acute phase of infection and 6-9 months later. (**C**) Ratio of ADCP to anti-spike IgG measured by S-Flow. (**D**) Schematic representation of complement-dependent cytotoxicity (CDC) assay. (**E**) Comparison of CDC activity in pre-pandemic samples, SARS-CoV-2-infected individuals in the acute phase of infection and 6-9 months later. (**F**) Ratio of CDC to anti-spike IgG as measured by S-Flow. (**G**) Schematic representation of antibody-dependent cellular cytotoxicity (ADCC). SARS-CoV-2 plasma induced NK degranulation as measured by CD107a staining using spike-expressing 293T cells as target cells. NK cells were isolated from healthy donors. (**H**) Comparison of ADCC activity in pre-pandemic samples, SARS-CoV-2-infected individuals in the acute phase of infection and 6-9 months later. (**I**) Ratio of ADCC to anti-spike IgG as measured by S-Flow. Statistical comparisons were performed by Mann Whitney test. The dashed line indicates the cutoff for positivity set based on values calculated following formula: cut-off = % mean positive cells from 19 pre-pandemic samples + 3x standard deviation. Each dot represents result from a single individual. Lines represent median and IQR.**p <0.01, ***p <0.001, and ****p < 0.0001. Pre-pandemic n=19, acute n=33, 6-9 months n=64.

Next, to evaluate the contribution of anti-S antibodies to complement dependent cytotoxicity (CDC), we assessed cell death in Raji cells engineered to express S protein in the presence of normal human serum as source of complement (Figure 2E, S5) (19). No differences in CDC activity was observed between the acute and late convalescent phase, where 60% and 56% of the subjects showed CDC activity, respectively (Figure 2F, G). The proportion of CDC-mediating antibodies within the total anti-S antibody fraction significantly increased between acute and late convalescence (p=0.0002, Figure 2H).

Killing of virus-infected cells can also be mediated by activated NK cells, after binding of immunocomplexes to CD16 (18). Therefore, antibody-dependent cellular cytotoxicity (ADCC) activity was measured using S-expressing 293T cells as target cells with degranulation measured by CD107a staining in primary NK cells as a readout for ADCC (Figure 2I, Figure S6). ADCC activity did not change between the acute and late convalescent phase (Figure 2J, K). At both time points, 59% - 66% of individuals showed anti-S mediated ADCC activity. However, similar to ADCP and CDC, the proportion of ADCC-mediating antibodies within the fraction of anti-S antibodies increased significantly over time (p<0.0001, Figure 2I). Analyzing only individuals with paired samples available, revealed similar results as the cohort as a whole (Figure S3, C-H). Overall, these data show that antibody effector functions mediated by S-specific antibodies remain stable over time and that the proportion of the functional antibody response within the total anti-S antibodies increases over time.

### SARS-COV-2 infection induces a sustained memory B cell compartment reacting against spike and nucleocapsid protein 6-9 months after infection

Upon re-infection, memory B cells are rapidly activated to differentiate into antibody-producing plasmablasts and/or re-initiate germinal centers in the case of secondary heterologous infection with antigenically similar pathogens (38). Therefore, they may play an important role in long-term immunity to SARS-CoV-2 and their evolving variants. We assessed the phenotype and frequency of antigen-specific memory B cells by staining with site-specific biotinylated recombinant S1 and N protein (Figure 3A, S7A, B). At late convalescence, 0.10% of the total CD27^+^ B cells are S1-specific, whereas 0.66% are N-specific (p<0.0001, Figure 3B). (Figure 3C, D). The proportion of CD27^+^CD38^+^ S1-specific B cells (75%, IQR=30%) is significantly increased compared to the proportion of CD27^+^CD38^+^ N-specific B cells (39%, IQR=26%, Mann-Whitney Test, p<0.0001) (Figure 3E). Moreover, the proportion of S1-versus N-specific B cells varies within each CD27^+^ B cell subset (p<0.0001, Figure 3F). We next analyzed S1- and N-specific B cells within the unswitched (IgD^-^IgM^+^) and switched (IgD^-^IgG^+^ and IgD^-^IgA^+^) B cell compartments (Figure S7A). S1-specific B cells were mainly IgD^-^IgG^+^, whereas N-specific B cells were either IgD^-^IgM^+^ or IgD^-^IgG^+^ (Figure 3G,H). The proportion of IgD^-^IgG^+^ S1-specific B cells (75%, IQR=24%) was significantly increased compared to the proportion of IgD^-^IgG^+^ N-specific B cells (37%, IQR=17%) (p<0.0001) (Figure 3I). Therefore, within each switched B cell subset, the proportion of S1-versus N-specific B cells was different (p<0.0001) (Figure 3J). Taken together, SARS-CoV-2 infection induces a robust memory B cell response targeting both S and N.

**Figure 3.**
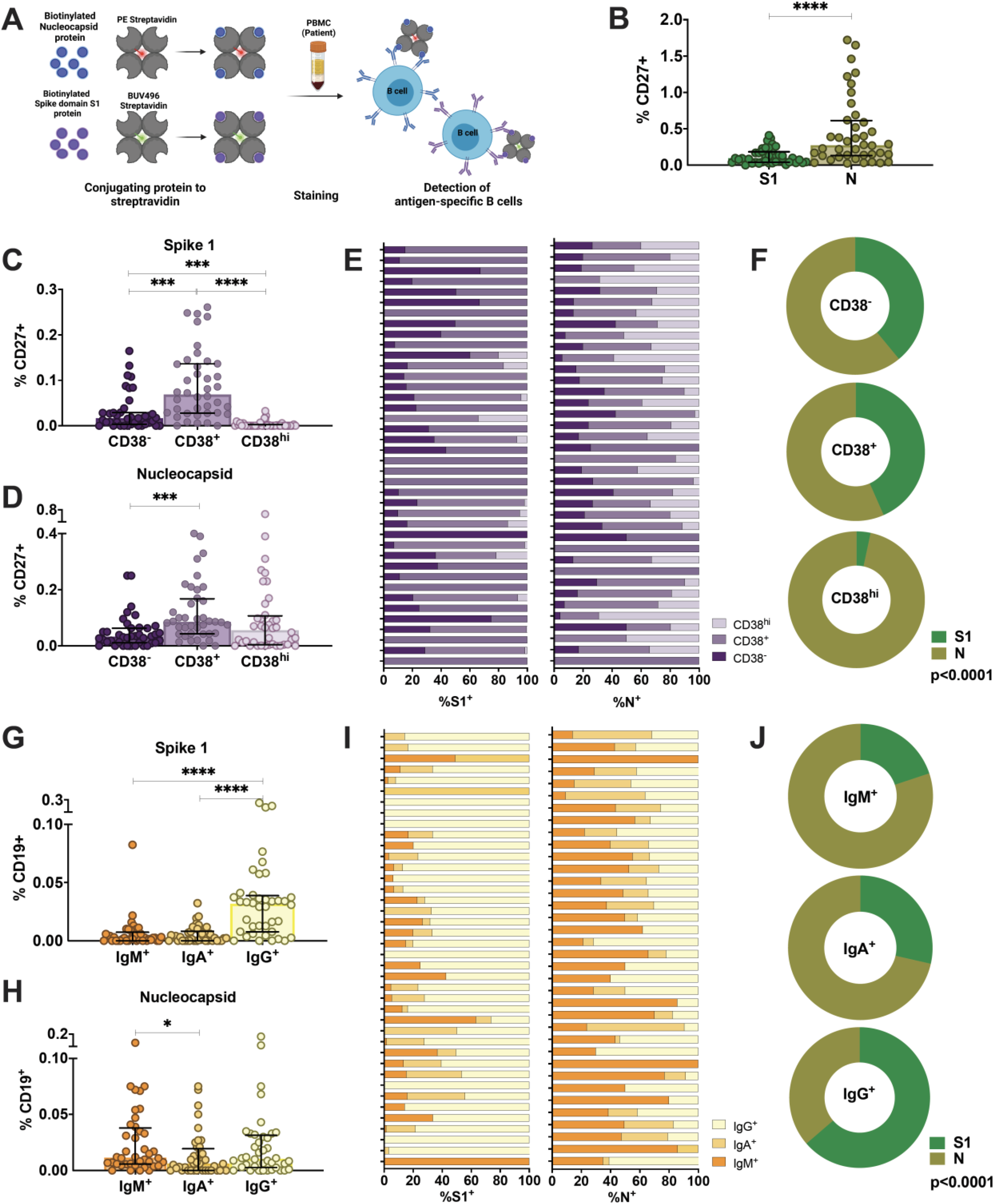
Characterization of antigen-specific memory B cells in the peripheral blood of individuals infected with SARS-CoV-2 6-9 months after infection. (**A**) Schematic representation of the memory B cell assay. (**B**) Comparison of percentages of S1-specific or N-specific memory B cells (CD19^+^CD27^+^). (**C, D**) Percentages of S1- and N-specific B cells among resting memory B cells (CD38^-^), activated memory B cells (CD38^+^) or plasmablasts within the CD27^+^ memory B cell population. (**E, F**) Proportion of S1-specific and N-specific CD27^+^CD19^+^ B cell subsets for each individual and the whole cohort (**G, H**) Percentages of S1- and N-specific cells in non-class-switched B cells (IgD^-^IgM^+^) or class-switched B cells (IgD^-^IgA^+^ or IgD^-^IgG^+^). (**I, J**) Proportion of S1-specific and N-specific switched and unswitched CD19^+^ B cells for each individual and for the whole cohort. Statistical comparisons were performed by Mann-Whitney test (B), Wilcoxon Rank Sum test (C, D, G, H) or Chi-square test (F, J). The dashed line indicates the cutoff for positivity set based on values calculated following formula: cut-off = % mean positive cells from 19 pre-pandemic samples + 3x standard deviation. Each dot represents result from a single individual (n=40). Lines represent median and IQR. **p <0.01, ***p <0.001, and ****p < 0.0001. n = 40. S1: subunit 1 of spike protein, N: Nucleocapsid protein.

### SARS-CoV-2 infection induces mainly spike and membrane protein-specific memory CD4^+^ and CD8^+^ T cells that are maintained up to 6-9 months after infection

In addition to humoral immunity, the generation and maintenance of virus-specific cellular immune responses is critical to help prevent reinfection. Long-term maintenance and phenotypes of SARS-CoV-2-specific memory T cell responses are still under investigation (24, 39, 40). SARS-CoV-2-specific CD4^+^ and CD8^+^ T cells were assessed in 33 individuals at late convalescence by incubating PBMCs with peptide pools covering immunodominant sequences of the viral S1, M and N protein (Figure 4A). Post incubation, activation induced marker (AIM) assays identified CD4^+^ antigen-specific cells using OX40^+^CD137^+^ combined with phenotypic markers to measure different memory and T helper (Th) subsets (Figure S8 A-D). Percentages of both S1- and M-specific CD4^+^ T cells were significantly increased compared to the percentage of N-specific cells (p<0.0001, p=0.0002), Figure 4B). Phenotypically, 42% of virus-specific T cells displayed an effector memory phenotype (CD45RA^-^CCR7^+^) and 87% of the cells showed a Th1-skewed phenotype (CXCR3^+^CCR6^-^) (Figure 4C, D). Comparing the memory phenotype of S1-, M- and N-specific cells, we observed that a lower proportion of S1-specific cells displayed an effector memory phenotype (23%) compared to M-specific cells (41%, p=0.0457) and N-specific cells (58%, p<0.0001) (Figure S9A). Moreover, 97% of M-specific cells showed a Th1-skewed phenotype compared to only 65% (p<0.0001) of the S1-specific cells and 71% (p=0.0130) of the N-specific cells (Figure S9B). In eight individuals, sufficient cell numbers were available to assess functionality by cytokine production after peptide stimulation using a multi-parameter *ex vivo* intracellular cytokine staining (ICS) assay (Figure S8E). SARS-CoV-2-specific CD4^+^ T cells produced Interleukin (IL)-2 (36%) or IL-6 (28%) after peptide stimulation, and were polyfunctional (Figure 4 E, F). Percentages of IL-2^+^ and IL-17^+^ cells were significantly higher after S1 stimulation compared to M stimulation (p=0.046, p=0.017) (Figure S9C).

**Figure 4.**
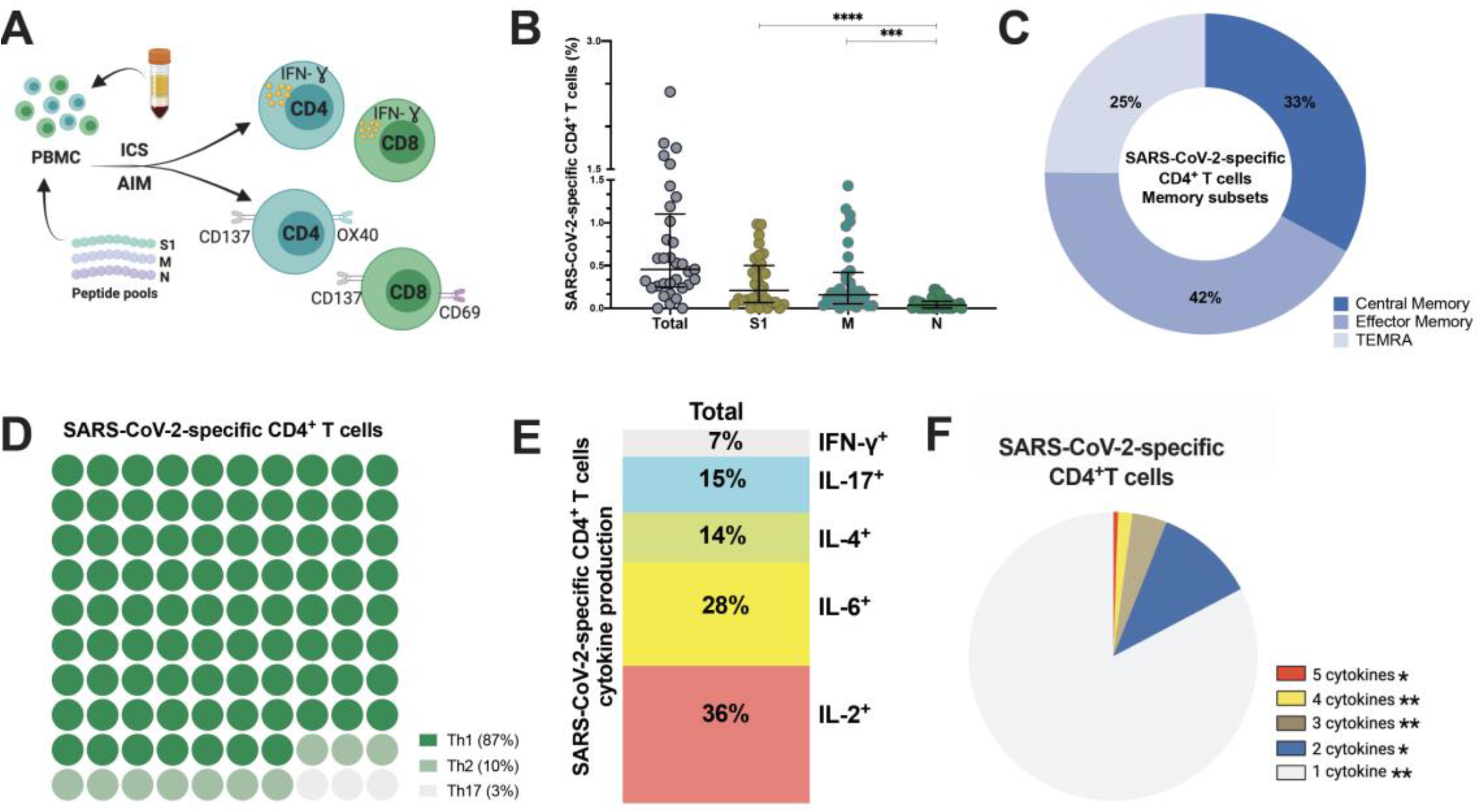
SARS-CoV-2-specific CD4^+^ T cells 6-9 months post-infection. **(A)** Schematic representation of the CD4^+^ T cell assay (**B**) Frequency (percentage of CD4^+^ T cells) of total SARS-CoV-2-specific CD4**^+^** T cells after overnight stimulation with S, M and N peptide pools as assessed by induced expression of OX40 and CD137. Each dot represents result from a single individual (n=8). Lines represent median and IQR, n=33 **(B)** Distribution of SARS-CoV-2– specific CD4**^+^** T cells among central memory, effector memory, and terminally differentiated effector memory cells (TEMRA). **(C)** Frequencies of SARS-CoV-2-specific CD4^+^ T helper (Th) subset. (**D)** Cytokine production and (**E**) pie chart representing the multifunctional SARS-CoV-2-specific CD4^+^ T cell response assessed by intracellular cytokine staining after incubation with SARS-CoV-2 peptides compared to unstimulated control. Statistical comparisons were performed by Kruskal-Wallis test (B) and Wilcoxon Rank Sum test (F). p< 0.05, **p < 0.01, ***p < 0.001, ****p<0.0001. S1: subunit 1 of spike protein; M: membrane protein; N: nucleocapsid protein.

Next, we assessed the frequency and phenotype of cytotoxic CD8^+^ T cells by AIM assay using CD69^+^CD137^+^ to identify antigen-specific CD8^+^ T cells. Frequency of total SARS-CoV- 2-specific CD8^+^ cells is 0.44% (Figure 5A) with 61% of these SARS-CoV-2-specific CD8^+^ T cells being terminally differentiated effector memory cells (TEMRA, CD45RA^+^CCR7^-^) (Figure 5B, Figure S9D). No differences were observed between S1-, M- and N-specific CD8^+^ T cells. Similar to antigen-specific CD4^+^ T cells, SARS-CoV-2-specific CD8^+^ T cells produced either IL-2 (56%) or IL-6 (16%) after peptide stimulation, and were polyfunctional (Figure 5 C, D), (Figure S9E). Interestingly, 2/33 (6%) individuals displayed no CD4^+^ T cell reactivity, and 6/33 (19%) individuals lacked a CD8^+^ T cell response after stimulation. In summary, sustained and functional CD4^+^ and CD8^+^ T cell responses are detected in the study participants, even after experiencing only mild or asymptomatic SARS-CoV-2 infection. These data suggest that SARS-CoV-2 can induce a long-lived cellular immune response, which could confer protection after reinfection or could be reactivated with vaccination.

**Figure 5.**
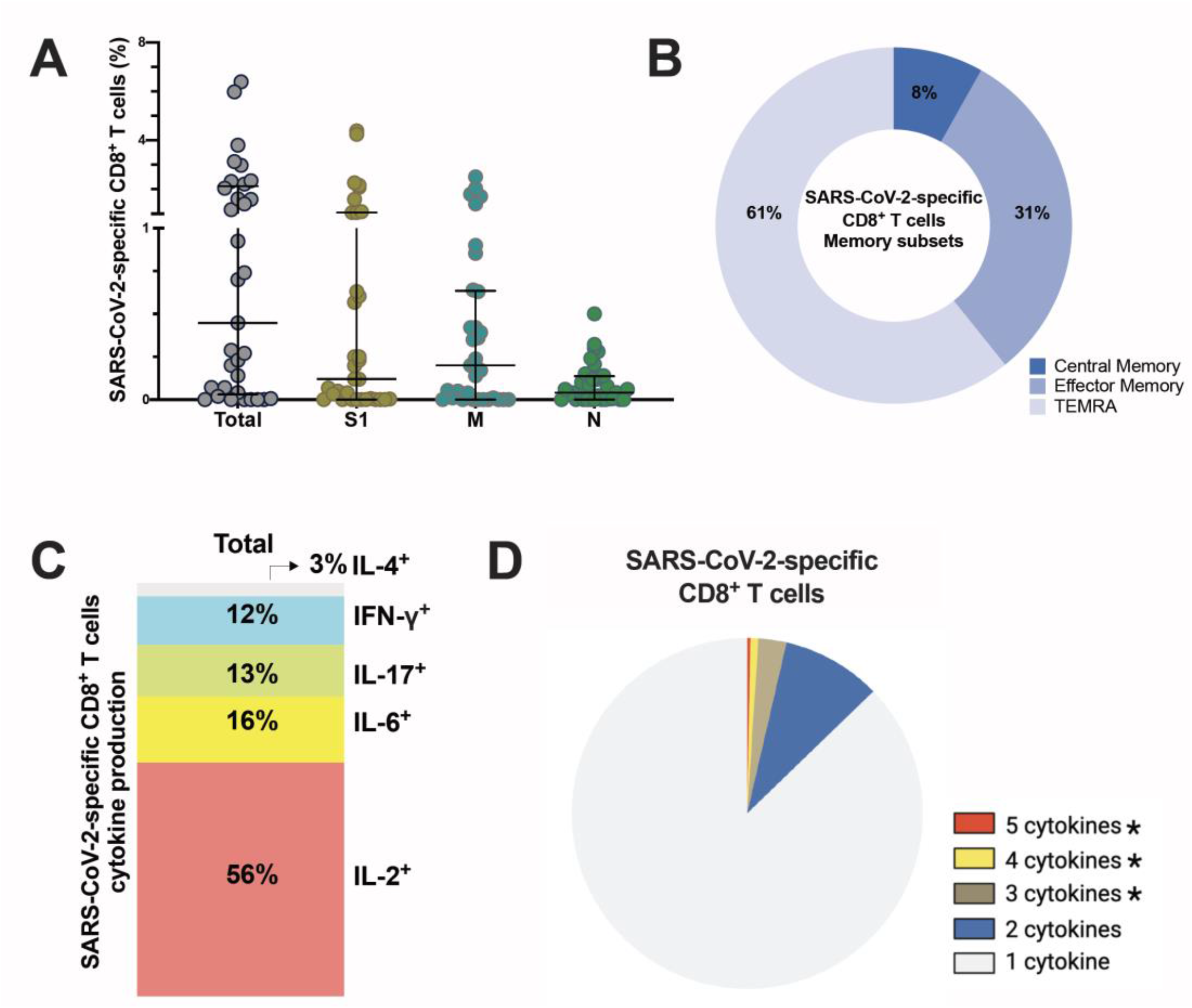
SARS-CoV-2-specific CD8^+^ T cells 6-9 months post-infection. **(A)** Frequency (percentage of CD8^+^ T cells) of total SARS-CoV-2-specific CD8**^+^** T cells after overnight stimulation with S, M and N peptide pools as assessed by induced expression of CD69 and CD137. Each dot represents result of a single individual (n=33). Lines represent median and IQR, n=33. **(B)** Distribution of SARS-CoV-2-specific CD8**^+^** T cells among central memory, effector memory, and terminally differentiated effector memory cells (TEMRA). **(C)** Cytokine production and **(D)** pie chart representing the multifunctional CD8**^+^** T of SARS-Cov-2-specific T cells assessed by intracellular cytokine staining after incubation with SARS-CoV-2 peptides compared to unstimulated control. Statistical comparisons were performed by **(A)** Kruskal-Wallis test and **(D)** Wilcoxon Rank Sum test. *p < 0.05, **p < 0.01, ***p < 0.001. S1: subunit 1 of spike protein; M: membrane protein; N: nucleocapsid protein.

### Symptomatic infection is associated with increased ADCC activity and increased frequency of SARS-CoV-2-specific CD4^+^ T cells observed 6-9 months after infection

In order to assess if symptomatic disease is associated to altered immune memory formation, we compared the functional immune response between asymptomatic and symptomatic patients with mild/moderate clinical presentation. Overall, no differences occurred in the humoral parameters assessed in the acute phase of infection (Figure 6A, S10). At late convalescence, symptomatic disease is associated with increased ADCC activity compared to asymptomatic individuals (p=0.0034) (Figure 6A). Moreover, percentages of N-specific CD27^+^ B cells, but not S1-specific, were increased in asymptomatic individuals versus patients who were symptomatic (p=0.051) (Figure 6B). Symptomatic disease was also significantly associated with increased percentage of SARS-CoV-2-specific CD4^+^ T cells (p=0.0018) with a central memory phenotype (p=0.0498) (Figure 6C, D). These data suggest that the outcome of acute infection has an imprint on the memory immune response with implications for response to subsequent infection or vaccination.

**Figure 6.**
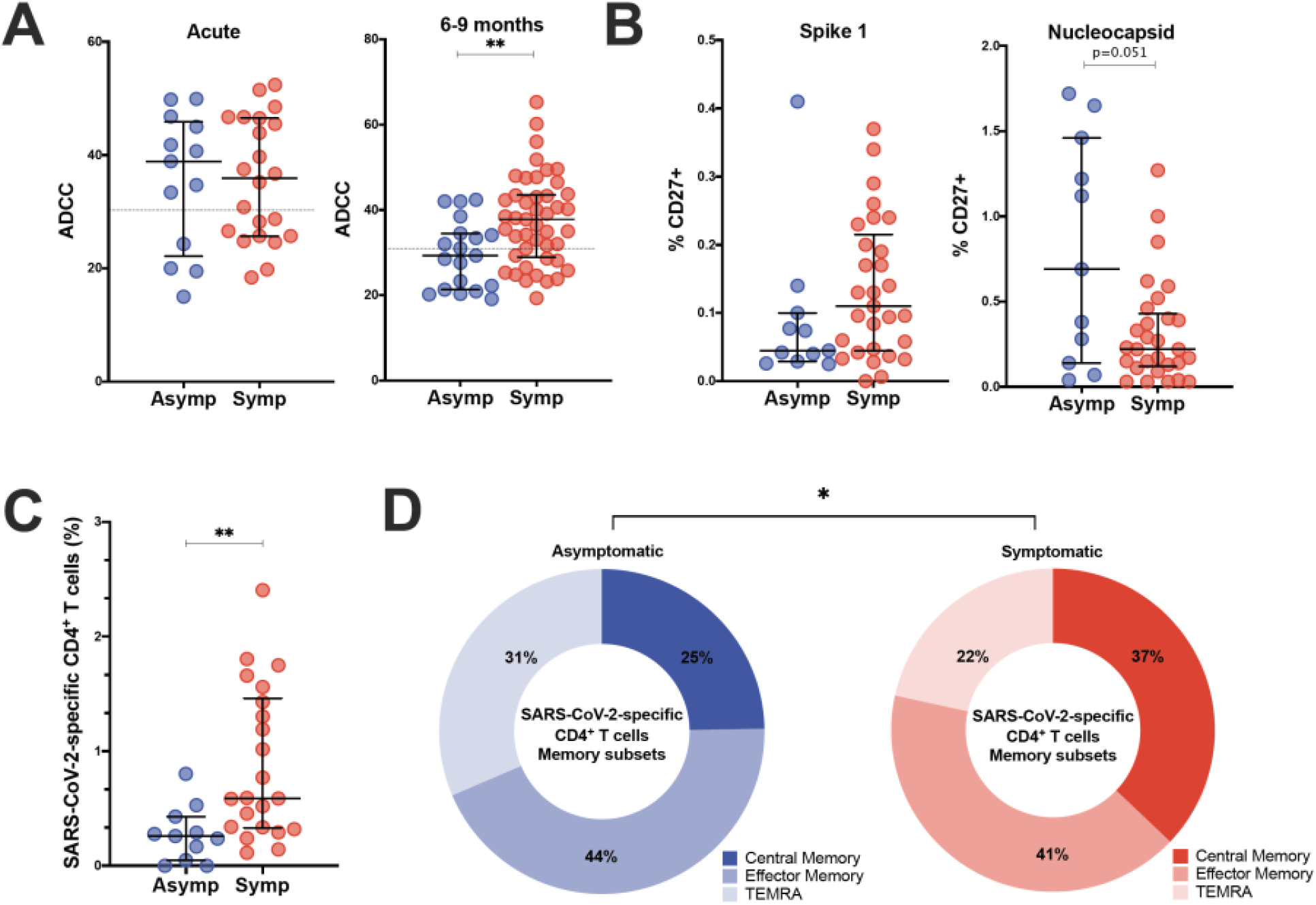
Comparison of adaptive immunity in asymptomatic and symptomatic individuals. (**A**). Comparisons of ADCC activity in asymptomatic (asymp; n=12) versus symptomatic (symp; n=20) individuals in the acute phase and 6-9 months after confirmed infection using 293T-spike cells as target cell. Percentage of CD107a positive cells is measured as readout for ADCC. **(B)** Comparison of percentages of S1-specific or N-specific memory B cells (CD19^+^CD27^+^) between 11 asymptomatic individuals and 29 symptomatic individuals**. (C)** Frequency (percentage of CD4^+^ T cells) of total SARS-CoV-2-specific CD4^+^ T cells after overnight stimulation with peptide pools comparing asymptomatic individuals (asymp; n=11) with symptomatic patients (symp; n=22). **(D)** Comparison of CD4^+^ T cell memory phenotype between asymptomatic individuals (asymp; n=11) and symptomatic patients (symp; n=22). Statistical comparisons were performed by (A, B and C) Mann Whitney tests and (D) Chi-square test for trend *p<0.05, **p <0.01.

### Correlations between various aspects of the functional anti-viral memory response

In order to assess the relation between antibody titers, functional humoral immunity, and the cellular T and B cell compartment we performed extensive correlation analysis (Figure 7). Age correlated to anti-S antibody titers and S1-specific CD19^+^IgD^-^IgG^+^ and CD19^+^CD27^+^B cells. In the acute phase of infection, anti-S IgG, IgM and IgA titers, functionality, measured by seroneutralization, and effector functions correlated. Seroneutralization, anti-S IgA, and ADCC correlated over time, albeit not very strong. Antibody effector functions 2-9 days post laboratory confirmation negatively correlated with total antigen-specific and S1-specific CD4^+^ T cells at late convalescence.

**Figure 7.**
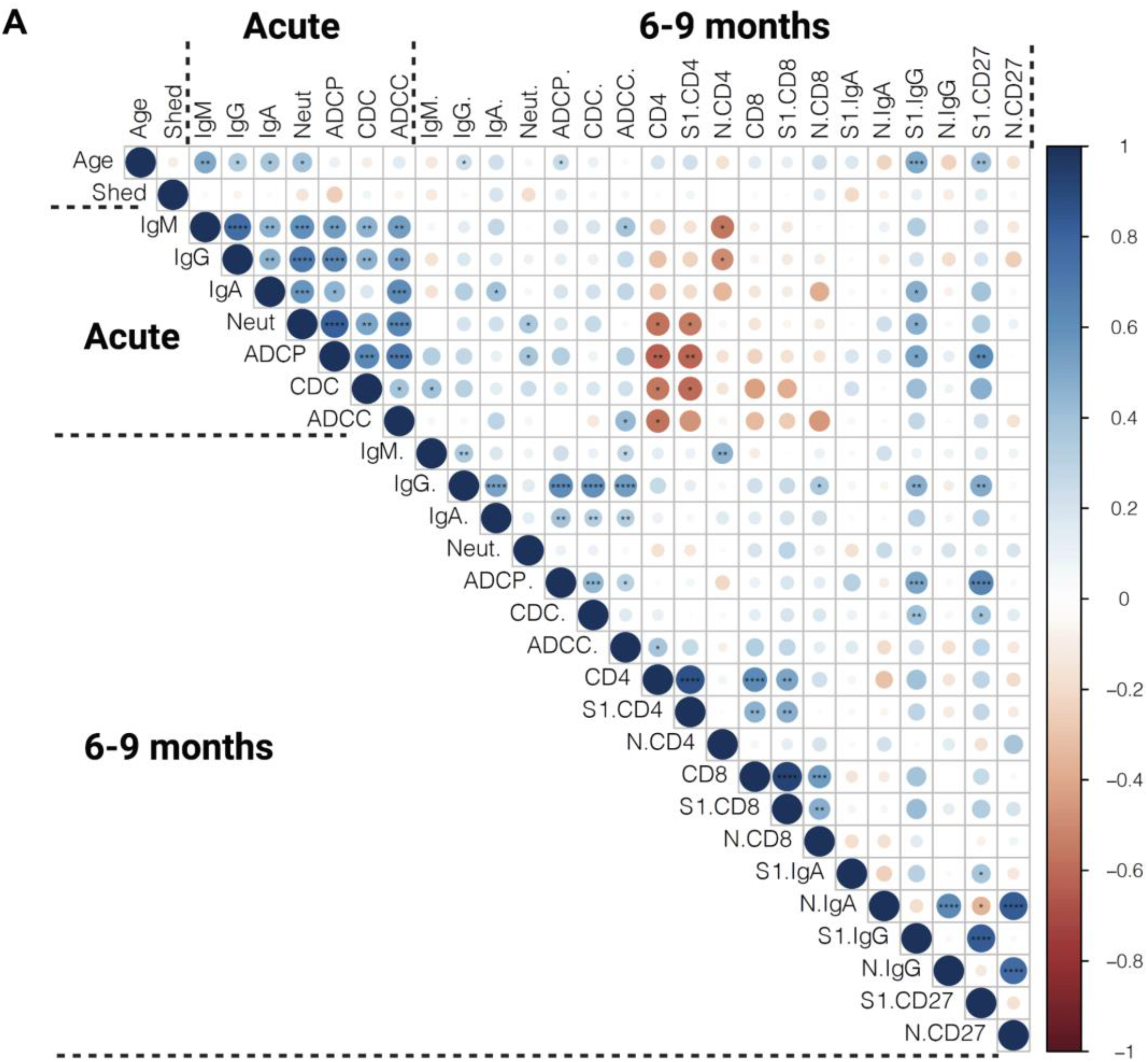
Spearman correlation matrix. Humoral immunity and effector functions measured in the acute phase and 6-9 months post infection were correlated to each other and to frequencies of antigen-specific B and T cells measured 6-9 months after infection. Red represents a negative correlation between two variables and blue indicates a positive correlation. The size of the dot represents the magnitude of the correlation coefficient. Statistical analysis was performed with spearman correlation test. *p <0.05, **p <0.01, ***p < 0.001, ****p < 0.001.

At late convalescence, anti-S IgG correlated with all three effector functions, but not with neutralizing capacity. Within the B cell compartment, N-specific IgG^+^, IgA^+^ and CD27^+^ B cells correlated to one another, as did S1-specific IgG^+^, IgA^+^ and CD27^+^ B cells. No correlation was identified between S1- and N-specific B cells. Anti-S IgG titers, ADCP, and CDC correlated with S1-specific IgG^+^ and CD27^+^ B cells. The S1-specific CD4^+^ T cell responses correlated with S1-specific CD8^+^ T cell responses, but did not correlate to antibody titers nor to effector functions or to S1-specific B cells. Overall, different aspects of a functional immune memory response do not fully correlate with one another and require separate evaluation when considering long-term immunity to SARS-CoV-2.

## Discussion

In this study, we investigated a partially asymptomatic cohort of Cambodian individuals in the acute and late convalescent phase for anti-S antibody titers, neutralization and effector functions, as well as SARS-CoV-2-specific B and T cell responses. As Cambodia was relatively COVID-19-free throughout 2020 (35), it is highly unlikely this cohort had additional exposure events after inclusion in this study, that could have boosted their immunity to SARS-CoV-2. One limitation is the uncertainty of the exact timing of exposure/infection, as infections were identified by screening at entry into Cambodia rather than in a direct surveillance or community cohort.

Studies assessing long-term immunity to SARS-CoV-2 in Asian populations are scarce (16, 32–34, 41). In addition, studies on cross-reactivity of the humoral and cellular compartment with other hCoVs have mainly focused on European and US populations (42–45). Historically, the population in East Asia seems to be more exposed to coronavirus-like viruses as only East Asian population show genetic adaptation to coronaviruses (46). The main natural reservoir of SARS-related coronaviruses is believed to be Horseshoe bats (genus *Rhinolophus*), which are endemic to Southeast Asia and China (47–49). Whether possible cross-reactivity to other coronavirus-like viruses or hCoVs may have influenced the adaptive immune response to SARS-CoV-2 in Southeast Asian populations remained to be investigated.

As expected, anti-S IgM, IgG and IgA titers declined over time and anti-S IgG becomes the major isotype at late convalescence (24, 50–52). In this study, IgA titers were the most affected over time. The formation of anti-S IgA is shown to be dependent on local lung inflammation (53–55) hence titers decline the strongest in asymptomatic/mild patients. Titers of neutralizing antibodies are reported to reach their maximum within the first month after infection and then decay, but mostly remain detectable six months and even up to one year after infection (10, 24). A relatively low rate of individuals retained neutralizing antibodies at late convalescence in this cohort (56%) as most longitudinal studies found 76-98% of individuals remaining positive (19, 24). This might be attributed to the absence of possible re-exposure and/or the consequence of asymptomatic/mild infection (10, 26, 39, 51). In contrast with other papers, neutralizing titers did not correlate to anti-S IgG antibodies at late convalescence. This might be due to varying relative contribution of anti-S IgG, IgM and IgA to SARS-CoV-2 neutralization at late convalescence, the genetic background of the participants or can be due to the different technique to measure anti-S binding and neutralizing antibodies (56, 57).

Fc-mediated effector functions contribute to clearance of virus-infected cells but are often critically overlooked. SARS-CoV-2 infection induces Fc-mediated effector functions irrespective of disease outcome (19–21). Antibody effector functions develop rapidly after infection and correlate with anti-S IgG and neutralizing titers in the acute phase and at late convalescence (19, 20). In this current study, between 55-66% of individuals showed antibody effector function activity up to nine months after infection. Also, ADCC persisted in a higher percentage of individuals compared to neutralization or other effector functions (22, 23). We report here the maintenance of CDC over time suggesting that both ADCC and CDC can contribute to protection from re-infection. ADCP levels decreased over time which could have consequences for antigen presentation and macrophage activation upon re-infection (58). Interestingly, the ratio of S-mediated effector functions over total anti-S IgG increases over time. Together with reports showing the evolution of the BCR repertoire over time (26, 59, 60), these data indicate ongoing affinity maturation and evolution of the antibody response to a more functional response. Therefore, measurement of only S-binding antibodies at late convalescence does not reflect their function.

S-, RBD- and N-specific memory B cells are maintained more than six months post symptom onset and their frequency increased over time (24, 61, 62). In this cohort, S1- and N-specific memory B cells persisted up to 6-9 months post infection with some variability between individuals. The percentage of S1-specific IgG B cells correlated with S-specific IgG antibodies, and S1-specific B cells displayed an activated phenotype. This suggests that these B cells could be recruited after secondary exposure with SARS-CoV-2 and might confer some level of protection against infection with new variants via additional diversification trough germinal center responses (38).

Anti-SARS-CoV-2 T cell immunity was assessed by AIM, which is a sensitive assay that provides a broader picture of the overall antigen-specific T cell response, compared to cytokine-detection based assays (6, 63). Persistence of functional memory T cells after SARS-CoV-2 infection has been reported, also after asymptomatic infection (24, 64). Similar to other reports, virus-specific memory CD4^+^ T cells were skewed to a Th1 or Th1/Th17 profile and displayed mainly an effector memory (CD45RA^-^CCR7^-^) phenotype (27, 64, 65). Virus-specific CD8^+^ T cells consisted mostly of cells with a TEMRA phenotype, a compartment of cytotoxic CD8^+^ T with limited proliferative potential (66). Polyfunctional virus-specific CD4^+^ and CD8^+^ T cells could be detected, mainly secreting IL-2 (16, 39), albeit we could only include few individuals in this analysis. Similar to other long-term cohorts, virus-specific CD4^+^ and CD8^+^ cells can be detected in up to 90% - 70% of the individuals, respectively (4).

Differences in frequency and phenotype of N- and S-specific B and T cells has been reported before (24, 27, 39, 67). This might be due to the difference in antigen availability, persistence, and immunological context. Together with other envelope proteins, S proteins cover the surface of the virus and bind to the host cell, while the N protein underlies viral packaging and hence is less accessible (68). The N protein is more conserved among coronaviruses (68), whereas S protein and especially the RBD-bearing S1 subunit are more prone for acquiring mutations (69, 70). Consequently, anti-N IgG rather than anti-S1 IgG can be found in individuals not exposed to SARS-CoV-2 (68, 71, 72). This might explain the observed higher frequency of N-specific B cells in our study.

Correlations between CD4^+^ T cells and humoral responses can be observed in some long-term cohorts (27, 29, 73, 74) but not all (24). In this study, there was no correlation between the S-specific cellular and humoral immune compartment at late convalescence. Therefore, neither anti-S IgG nor neutralizing antibodies are a good proxy to determine the cellular response to SARS-CoV-2. Moreover, correlations between anti-S antibody titers and Fc-related functions at late convalescence are weak and subtle differences might lead to a different disease outcome upon re-exposure. Hence, serological testing alone might not be sufficient to understand the full spectrum of long-term immunity generated after SARS-CoV-2 infection.

The development, characteristics and functionality of the totality of long-term immunity in asymptomatic infected individuals remains to be further characterized. We observed an increase of ADCC at late convalescence in patients who had mild/moderate disease compared to asymptomatic individuals. This observation is in line with studies showing increased anti-S IgG afucosylation in severe patients compared to mild and asymptomatic cases (75, 76). Indeed, afucosylated monoclonal antibodies can cause elevated ADCC though increased IgG-FcγRIIIa affinity (77, 78). More severe COVID-19 induced a stronger SARS-CoV-2-specific CD4^+^ T cell response (27–29). We confirm and extend these data as we observed lower levels of virus-specific CD4^+^ T cells in asymptomatic individuals compared to mild/moderate cases. These data suggest that different disease outcome after infection results in altered long-term immunity, which could shape the response to subsequent infection or vaccination.

Taken together, our work shows additional evidence of long-term and persistent immunity after asymptomatic and mild SARS-CoV-2 infection. Furthermore, this cohort describes the immune response in individuals of Asian origin and in the absence of re-exposure to SARS-CoV-2. We show the persistence of humoral immunity, antibody effector functions, and virus-specific memory T and B cells 6-9 months after infection, which do not correlate to each other. These data enhance our understanding of long-term functional immunity.

## Methods

### Study population

Ethical approval for the study was obtained from the National Ethics Committee of Health Research of Cambodia. Written informed consent was obtained from all participants prior to inclusion in the study. Pre-pandemic blood samples were obtained from clinically healthy individuals included in the dengue vaccine initiative study in 2015-2016. Clinically healthy adult volunteers who presented at the International Vaccination Centre, Institut Pasteur du Cambodge before the onset of the pandemic were included to validate the antigen-specific B and T cell staining. Acute SARS-CoV-2 infected patients were identified via screening of imported cases in Cambodia between 6^th^ March to 12^th^ August 2020. All laboratory confirmed cases are quarantined and monitored for symptoms. Moreover, 1-15 follow-up nasopharyngeal/oropharyngeal swab samplings for SARS-CoV-2 detection were conducted to assess viremia. Patients were only discharged after two consecutive negative RT-PCR tests within 48h. Symptomatic patients displayed mild/moderate symptoms such as running nose, cough, fever and difficult to breath. In total, we included 64 individuals for follow up. In 33 individuals, 2-9 days after laboratory confirmation, a blood sample was obtained. A second blood sample was obtained 6-9 months later from all 64 study participants. Participant characteristics and clinical signs are summarized in Table S1. Plasma was collected and stored at −80°C, The PBMCs were isolated via Ficoll-Paque separation, cryopreserved in 10% DMSO/FBS and stored in liquid nitrogen until analysis. The National Institute for Biological Standards and Control (NIBSC) 20/130 (research reagent) and 20/118 (reference panel) have been obtained from WHO Solidarity II, the global serologic study for COVID-19.

### SARS-CoV-2 detection

Molecular detection of SARS-CoV-2 in combined nasopharyngeal/oropharyngeal swabs was performed as previously described (36). Briefly, RNA was extracted with the QIAamp Viral RNA Mini Kit (Qiagen) and real-time RT-PCR assays for SARS-CoV-2 RNA detection were performed in using primers/probes from Charité Virologie (Berlin, Germany (79)) to detect both E and RdRp genes.

### Virus neutralization assay

The detection of neutralizing antibodies was achieved by foci reduction neutralization test (FRNT) similar as described before (80) and adapted to SARS-CoV-2. Briefly, serial diluted, heat-treated plasma samples were incubated with a Cambodian SARS-CoV-2 isolate (ancestral strain; GISAID: EPI_ISL_956384; (36)) for 30min at 37°C and 5% CO_2_. The mixtures were distributed on African green monkey kidney cells (VeroE6; ATCC CRL-1586) and incubated again for 30min 37°C and 5% CO_2_. Afterwards, the mixtures were replaced by an overlay medium containing 2% carboxymethyl cellulose (Sigma-Aldrich) in Dulbecco’s modified Eagle medium (DMEM; Sigma-Aldrich) supplemented with 3% FBS (Gibco) and 100 U/mL penicillin-streptomycin (Gibco). Infection was visualized 16-18h after inoculation by staining of infected cells with a SARS-CoV-2-specific antibody (rabbit, antibodies-online GmbH), targeting the S2 subunit of the viral spike protein, and afterwards with antibody anti-rabbit IgG HRP conjugate (goat; antibodies-online GmbH). Finally, cells were incubated with TrueBlue TMB substrate (KPL), and infection events appear as stained foci and were counted with an ELISPOT reader (AID Autoimmune Diagnostika GmbH, Strassberg, Germany). The amount of neutralizing antibodies is expressed as the reciprocal serum dilution that induces 50% reduction of infection (FRNT50) compared to the positive control (virus only) and is calculated by log probit regression analysis (SPSS for Windows, Version 16.0, SPSS Inc., Chicago, IL, USA). FRNT50 titers below 10 are considered negative.

### S-expressing cell lines

Transfected cell lines, Raji (ATCC® CCL-86™) and 293T (ATCC® CRL-3216™), with SARS-Cov-2 spike plasmid or a control plasmid using Lipofectamine 2000 (Life technologies) are kind gifts from Olivier Schwartz, Institut Pasteur, Paris, France (19). Spike-expressing Raji cells and Raji control cells were cultured at 37°C, 5% CO2 in RPMI medium while 293T-spike cells and 293T control cells were cultured in DMEM medium. All media were completed with 10% FBS (Gibco, MT, USA), 1% L glutamine (Gibco), 1% penicillin/streptomycin and puromycin (1 μg/mL, Gibo™) for cell selection during the culture.

### S-Flow assay

The S-Flow assay was performed as previously described (37). Briefly, plasma samples were diluted (1:200) in 1xPBS with 2mM EDTA and 0.5% BSA (PBS/BSA/EDTA) and incubated with 293T-spike cells (80000 cells/100µl) for 30 minutes on ice. The cells were washed with PBS/BSA/EDTA and stained either with anti-IgM PE (dilution 1:100, Biolegend) and anti-IgG Alexa Fluor^TM^ 647 (dilution 1:600, Thermo Fisher) or anti-IgA Alexa Fluor 647 (dilution 1:800, Jakson ImmunoResearch) for 30 minutes on ice. The cells were washed with 1xPBS and fixed using buffer of the True-Nuclear Transcription Factor Staining kit (Biolegend). After fixing, the cells were washed and resuspended in 1xPBS. The results were acquired using FACS Canto II, BD Biosciences. The gating strategy for anti-IgM, anti-IgG or anti-IgA positive cells was based on the 293T control cells incubated with negative SARS-CoV-2 reference plasma. The data were reported as percentage of positive cells for anti-IgM, anti-IgG or anti-IgA. The NIBSC Research Reagent (20/130) and panel (20/118) (WHO Solidarity II) was utilized to set the cutoff for positivity based on the background staining of the negative SARS-CoV-2 plasma and calculated following formula: cut-off= % positive cells + 2x standard deviation.

### Antibody dependent cellular phagocytosis (ADCP) assay

THP-1 cells (ATCC® TIB-202^TM^) were used as phagocytic cells. For this, 1 µg of biotinylated S1 protein (Genscripts) was used to saturate the binding sites on 1 µl of FluoroSphere neutravidin beads (Thermo Fisher) overnight at 4°C. Excess protein was removed by washing the pelleted beads. The protein-coated beads were incubated with 40 µl heated-inactivated plasma diluted in complete RPMI (1:40) for 15 minutes at room temperature. Then, 5×10^4^ THP-1 cells suspended in 50 µl complete RPMI were added to the complex and incubated for 16 hours at 37°C, 5% CO_2_. After incubation, the cells were washed with 1xPBS and fixed using buffer in True-Nuclear Transcription Factor Staining kit (Biolegend). After fixing, the cells were washed and resuspended in 1xPBS. The samples were analyzed using FACS Canto II, BD Biosciences. Phagocytosis activity was scored by the integrate mean fluorescence intensity (iMFI) value (% positive fluorescence THP-1 cells x MFI of the positive fluorescence THP-1 cells).

### Complement dependent cytotoxicity (CDC) assay

The assay used spike-expressing Raji cells as target cells, pooled serum (4 healthy donors) as complement source and heated-inactivated patient plasma as antibody source. In short, 50 µl of heated-inactivated plasma (1:50) were incubated with Raji-spike cells for 30 minutes at 37°C, 5% CO_2_. Afterward, 50 µl of complete RPMI containing 15% of pooled serum was added into the cells and incubated at 37°C, 5% CO_2_ for 14 hours. The cells were washed with PBS and stained with Zombie Aqua viability dye (BioLegend) for 20 minutes on ice and then stained anti-APC C3/C3b/iC3b antibody (Cedarlane) for 30 minutes on ice. The cells were fixed with fixation buffer in True-Nuclear Transcription Factor Staining kit (Biolegend) for 20 minutes on ice. After fixing, the cells were washed and resuspended in 1xPBS. The samples were acquired using FACS Canto II, BD Biosciences. The results were reported as percentage of cell death and MFI of C3 deposition on the cells.

### Antibody-dependent cellular cytotoxicity (ADCC) assay

The assay used 293T-spike cells as a target cell and purified NK cells from healthy donor PMBCs as effector cells. First, 293T-spike cells were incubated with heated-inactivated patient plasma diluted in complete DMEM medium (1:50) at 37°C, 5% CO_2_ for 30 minutes. The NK cells were enriched by magnetic negative selection (Miltenyi) according to manufactor’s instruction. The 293T-spike cells were washed five times with complete RPMI medium. The NK cells were mixed with 293T-spike cells at a ratio 1:1 at final volume of 100 µl complete RPMI. Anti-CD107a and Monensin (Biolegend) 1:1000 dilution were added to the suspension and incubated at 37°C, 5% CO_2_ for 6 hours. The cells were washed with 1xPBS and stained with Zombie Aqua viability dye (BioLegend) for 20 minutes on ice. Then the cells were stained with anti-CD3 and anti-CD56 for 30 minutes on ice. The cells were washed and fixed/permeabilized using True-Nuclear Transcription Factor Staining kit (Biolegend) for 20 minutes on ice. After staining, the cells were washed and resuspended in 1xPBS. The samples were acquired using FACS Canto II, BD Biosciences.

### Detection of antigen-specific memory B cells

Biotinylated SARS-CoV-2 S1 protein and biotinylated SARS-CoV-2 N protein were purchased from GenScript. The biotinylated proteins were combined with different streptavidin (SA) fluorophore conjugates, BUV496 (BD Biosciences) and PE (Biolegend), respectively, at 1:1 molar ratio. Briefly, each SA was added gradually (3 times, every 20 minutes) to 20 µl of each biotinylated protein (1 µM) on ice. The reaction was quenched with D-biotin (GeneCopeia) at 50:1 molar ratio to SA for a total probe volume of 30 µl for 30 minutes on ice. Probes were then used immediately for staining. Each staining used 5 µl of probe. Shortly, patient PBMCs was washed with 1xPBS and stained with Zombie Aqua viability dye (BioLegend) for 10 minutes on ice. The cells were stained with the probes. Then the cells were washed and stained with anti-IgG antibody, for 30 minutes on ice. After that, the cells were washed and stained with master mix containing of anti-CD3, anti-CD19, anti-CD27, anti-CD38, anti-IgD, anti-IgM and anti-IgA antibodies for 30 minutes on ice Antibodies are listed in Table S2. After staining, the cells were washed and resuspended in 1xPBS with 2% FBS. The samples were analyzed using FACS Aria, BD Biosciences. The flow cytometry gating strategy to classify memory B cell subsets and switched B cells is shown in Figure S7. Overall, 40 samples were of sufficient quality and were included in the analysis.

### Activation-induced markers (AIM) T cell assay

Antigen-specific CD4^+^ and CD8^+^ T cells, as well as memory T cells and T helper subsets were assessed by Activation-Induced Marker (AIM) assay (6, 24). Cells were cultured at 37°C, 5% CO_2_, in the presence of SARS-CoV-2-specific S1, M and N protein pools [1 µg/mL] (PepTivator® SARS-CoV-2 regents; Miltenyi Biotec) in 96-well U-bottom plates at 0,5-1×10^6^ PBMCs per well. After 24 hours, cells were washed in 1xPBS supplemented with 0.5% bovine serum albumin (BSA) and 2 mM EDTA (FACS buffer) and stained with Zombie Aqua Fixable Viability kit (Biolegend) and incubated for 20 min at 4°C followed by surface staining for 30 min at 4°C. Stained cells were washed and resuspended in FACS buffer and analyzed using a FACSAria Fusion (BD Biosciences). Antibodies are listed in Table S2. Negative controls without peptide stimulation were included for each donor. Antigen-specific CD4^+^ and CD8^+^ T cells were measured subtracting the background (unstimulated control) from the peptide-stimulated sample. Negative results were set to zero. Data were analyzed with FlowJo software version 10.7.1 (FlowJo LLC). Overall, 33 samples were of sufficient quality and were included in the analysis.

### Intracellular staining (ICS) assay

Functional SARS-CoV-2-specific CD4^+^ and CD8^+^ T cells were assessed by surface and intracellular staining in a subset of individuals if sufficient amount of PBMCs were obtained (n=8). Cells were cultured at 37°C, 5% CO2, in the presence of SARS-CoV-2-specific S1, M and N protein pools separately [1 µg/mL each] (PepTivator® SARS-CoV-2 reagents; Miltenyi Biotec), Monensin (Biolegend) 1:1000 dilution and anti-Human CD28/CD49d purified [100 µg/mL] (BD Bioscience) in 96-well U-bottom plates at 0,5-1×10^6^ PBMCs per well. After 6 hours, cells were washed in FACS buffer and stained using a Zombie Aqua Fixable Viability kit (Biolegend) and incubated for 20 minutes at 4°C. Cells were then washed in PBS and fixed/permeabilized with True-Nuclear™ Transcription Factor Buffer Set (Biolegend). Surface (CD3, CD4 and CD8) and intracellular markers (IFN-γ, IL-2, IL-4, IL-6 and IL-17) were detected via the subsequent addition of directly conjugated antibodies incubating for 30 minutes at 4°C. Antibodies are listed in Table S2. Stained cells were finally washed and resuspended in FACS buffer and analyzed using a FACSAria Fusion (BD Biosciences). Antigen-specific CD4^+^ and CD8^+^ T cells were measured subtracting the background (unstimulated control) from the peptide-stimulated sample. Negative results were set to zero. Data were analyzed with FlowJo software version 10.7.1 (FlowJo LLC).

### Statistical analysis

Calculations, figures and statistics were made using Prism 9 (GraphPad Software) or RStudio (Version 1.2.1335). The data were tested for statistical normality before applying the appropriate statistical tests. All information about sample sizes and statistical tests performed were shown in the figure legends. Spearman correlation plot was calculated and visualized with the following packages: FactoMineR, factoextra (https://cran.r-project.org/web/ packages/factoextra/index.html) and corrplot (https://github.com/taiyun/corrplot) in R (Version 3.6.1) and RStudio (Version 1.2.1335).

### Data availability

All data associated with this study are available in the main text or the supplementary materials.

## Acknowledgements

This publication has been supported by WHO Solidarity II, global serologic study for COVID-19, with funding from the German Federal Ministry of Health (BMG) COVID-19 Research and development to WHO. We would like to acknowledge all patients that participated to the study. We would like to thank Borita Heng for her technical assistance.

The study was funded by the Howard Hughes Medical Institute (HHMI)–Wellcome Trust (208710/Z/17/Z to T.C.), and « URGENCE COVID-19 » fundraising campaign of Institut Pasteur (T.C., H.A.,P.D.). H.A. is supported by the German Centre for International Migration and Development (CIM). The graphical abstract was created with BioRender.com

## Author contributions

Conceptualization: TC, PD, EAK; Methodology: HV, AM, HA, LS, SS, NY, PP; Investigation: HV, AM, HA, BT, DV, TC, EAK; Visualisation: HV, AM, HA, SL; Funding acquisition: TC, HA, PD; Patient inclusion: HS, SS; Cohort management and patient selection: SL, SL; Project administration: TC, EAK, PP, PD; Supervision: HV, AM, BT, SO, DV, EAK, TC; Writing, original draft: HV, AM, HA, TB, EAK, TC; Writing, review and editing: HV, AM, HA, TB, VD, OS, EAK, TC

## Supplemental information

### Supplementary Material

**Figure S1.**
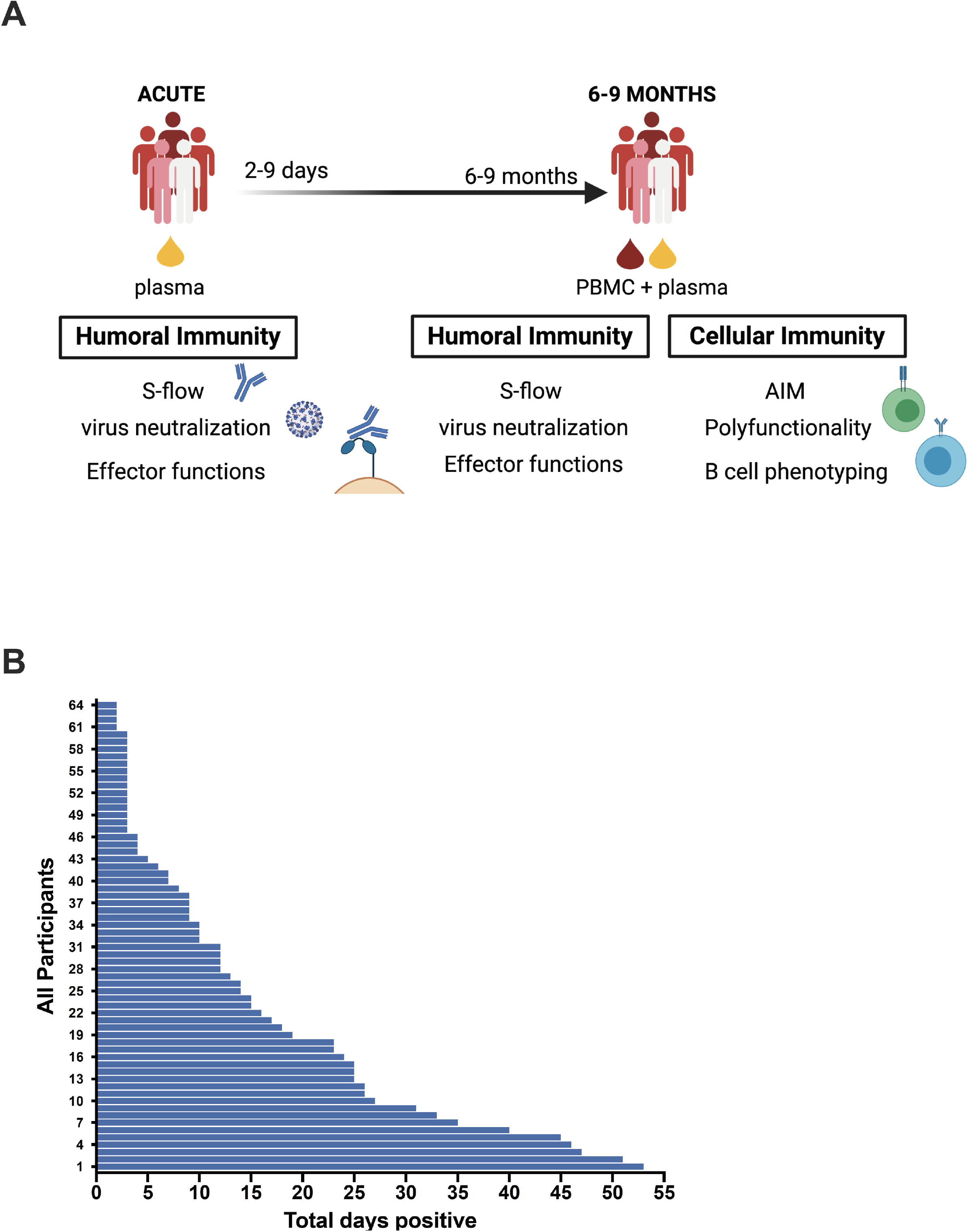
Characterization of the cohort. (**A**) Graphical summary of the assays performed on samples of the different time points. (**B**) Length of RNA shedding in the patients.

**Figure S2.**
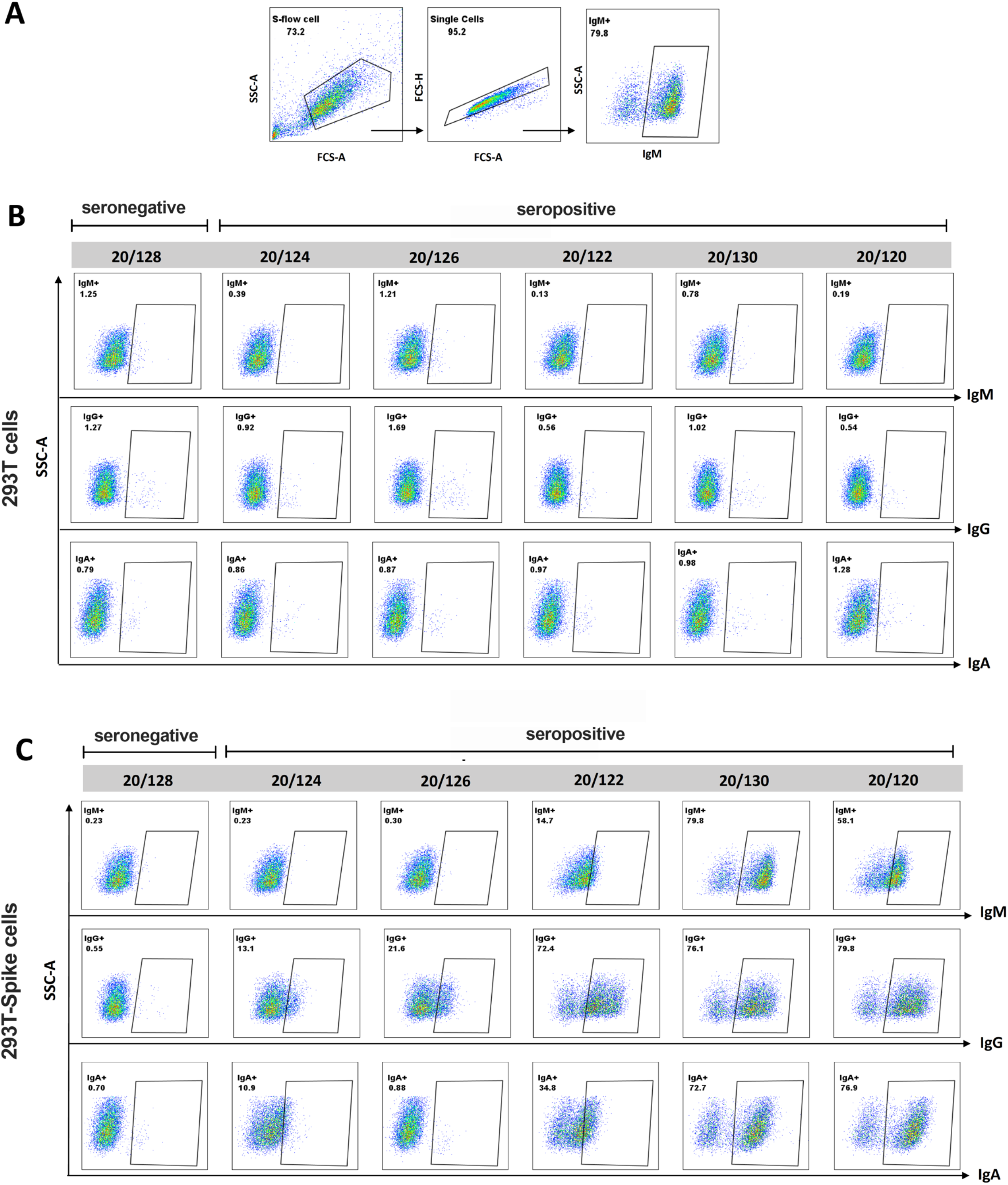
Measurement of anti-S-binding antibodies using S-Flow. (**A**) Representative gating strategy (**B-C**) Percentages of spike-binding Ig were measured in plasma as percentage of cells, which are positive for anti-IgM, anti-IgG or anti-IgA. Specific binding was calculated as 100 x (% binding on S-expressing 293T cells - % binding on 293T control cells)/(100-% of binding to 293T control cells). Graph shows data from triplicate testing of the COVID-19 reference plasma panel (NIBSC 20/120, 20/124, 20/124, 20/126, 20/128, 20/130) obtained from the NIBSC.

**Figure S3.**
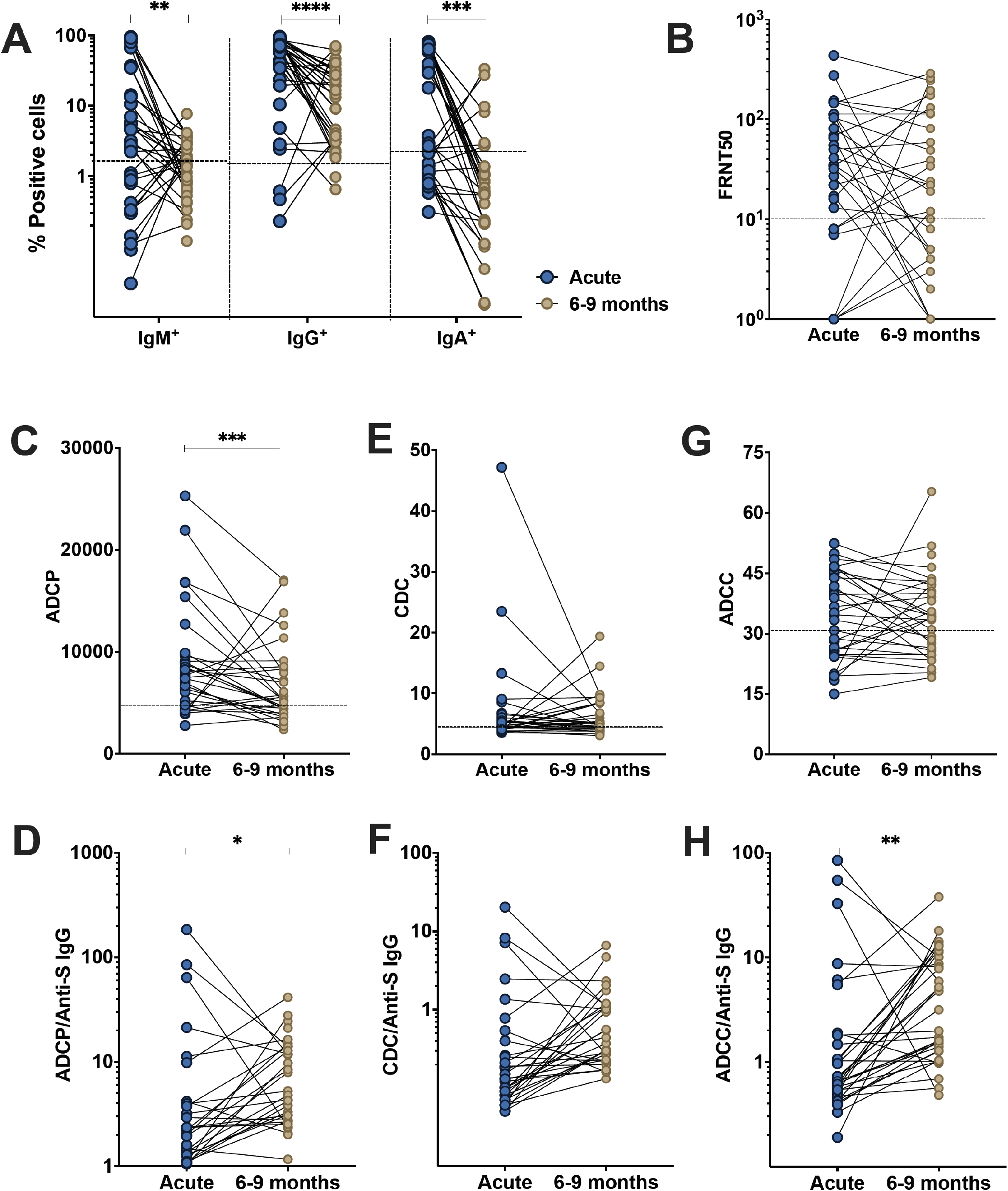
Analysis of antibody features in paired patient samples. Individuals were sampled 2-9 days post laboratory confirmation and 6-9 months later. (**A**) Amount of antibodies against spike protein were reported as percentage of spike-expressing 293T cells bound by IgM, IgG, IgA in the S-Flow assay. (**B**) SARS-CoV-2 neutralizing activity was calculated as FRNT50 titer in foci reduction neutralization test. (**C**) Comparison of ADCP activity in SARS-CoV-2-infected individuals in the acute phase of infection and 6-9 months post-infection. (**D**) Ratio of ADCP to anti-spike IgG measured by S-Flow. (**E**) Comparison of CDC activity in SARS-CoV-2-infected individuals in the acute phase of infection and 6-9 months post infection. (**F**) Ratio of CDC to anti-spike IgG measured by S-Flow. (**G**) Comparison of ADCC activity in SARS-CoV-2-infected individuals in the acute phase of infection and 6-9 months post infection. (**H**) Ratio of ADCC to anti-spike IgG measured by S-Flow. Statistical comparisons were performed by Wilcoxon test. The dashed line indicates the cutoff for positivity based on values calculated following formula: cut-off = % mean positive cells from 19 pre-pandemic samples + 3x standard deviation. Each dot represents result from a single individual. Lines represent median and IQR. Each dot represents one individual. Lines represent median and IQR. *p < 0.05, **p <0.01, ***p <0.001, and ****p < 0.0001. IgM/IgA/IgG antibody: n=33, Neutralization: n= 33, ADCP: n=30, CDC: n=30, ADCC n=32.

**Figure S4.**
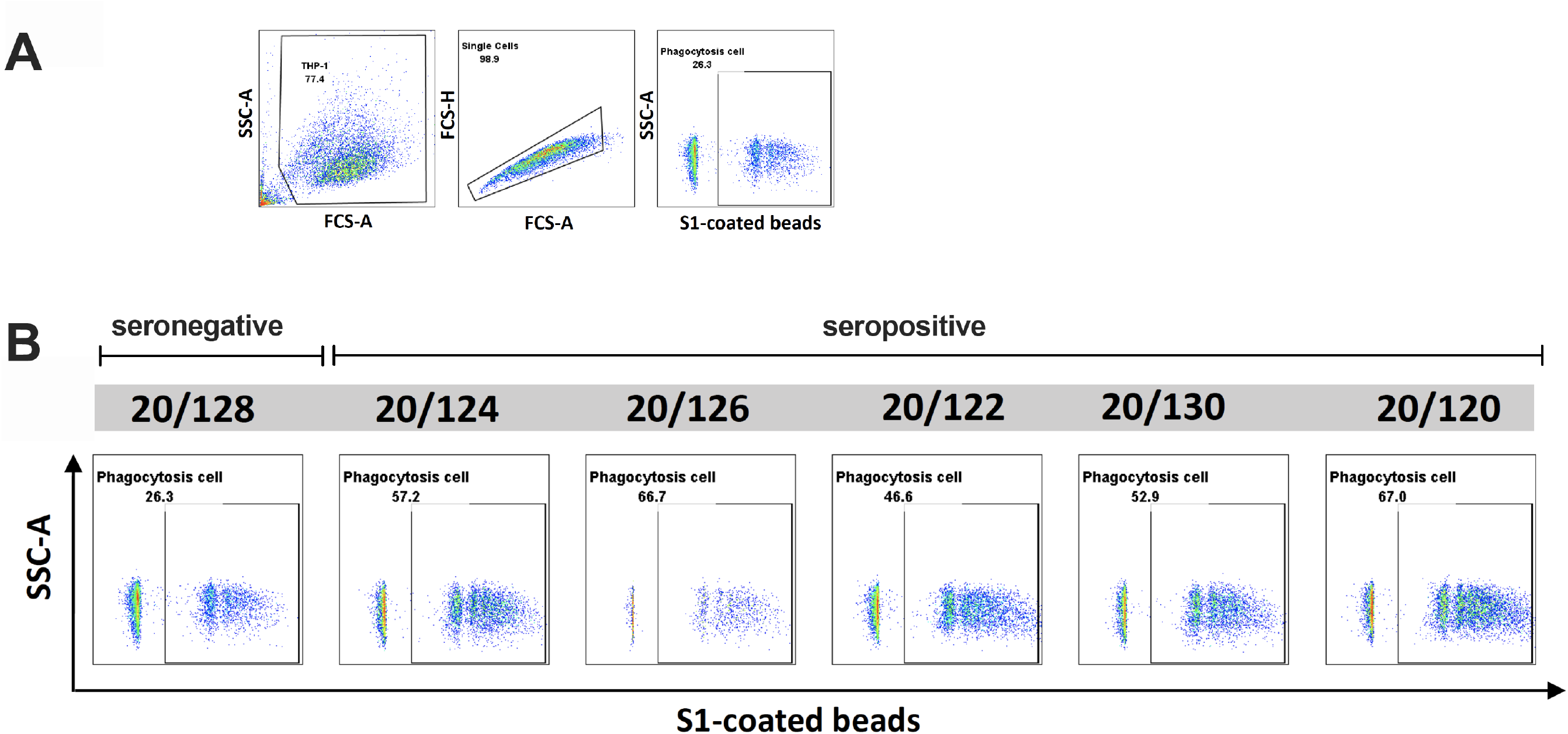
Antibody-dependent cellular phagocytosis (ADCP) assay. Quantification of the S1-coated beads engulfed by THP-1 cells. (**A**) Representative gating strategy. (**B**) ADCP was defined by the percentages of THP-1 cells which are positive for FITC-neutravidin beads coated with biotinylated S1 protein. Representative ADCP assay using the COVID-19 reference plasma panel (NIBSC 20/120, 20/124, 20/124, 20/126, 20/128, 20/130) obtained from the NIBSC. The ADCP activity represents the integrate mean fluorescence intensity (iMFI) value (% positive fluorescence THP-1 cells x MFI of the positive fluorescence THP-1 cells). The experiment was performed in duplicate.

**Figure S5.**
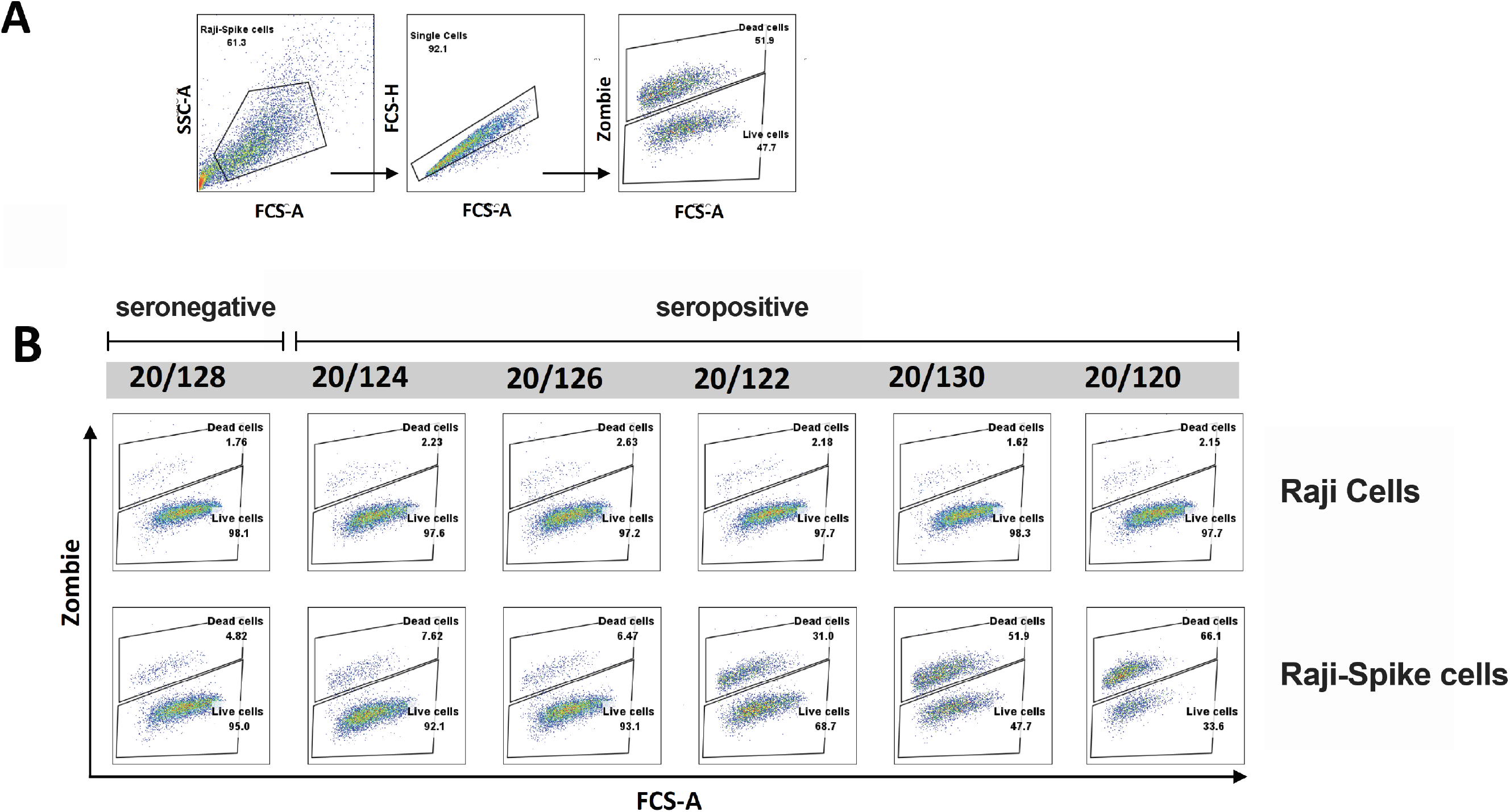
Complement-dependent cytotoxicity (CDC) assay. SARS-CoV-2 plasma induces C3 deposition and cell death in spike-expressing Raji cells. (**A**) Representative gating strategy. (**B**) CDC assay was performed by using pike-expressing Raji cells as target cells, serum (pooled from 3 healthy donors) as complement source and heat-inactivated plasma from SARS-CoV-2 patients or from reference panel as antibody source. CDC activity was measured as the percentage of C3^+^Zombie^+^ pike-expressing Raji cells after incubation with plasma and complement. Representative CDC assay using the COVID-19 reference plasma panel (NIBSC 20/120, 20/124, 20/124, 20/126, 20/128, 20/130) obtained from the NIBSC. Complement-induced cell death was calculated as percentage of C3^+^ dead cells of total cells using spike-expressing Raji cells as target cell. The experiment was performed in duplicate.

**Figure S6.**
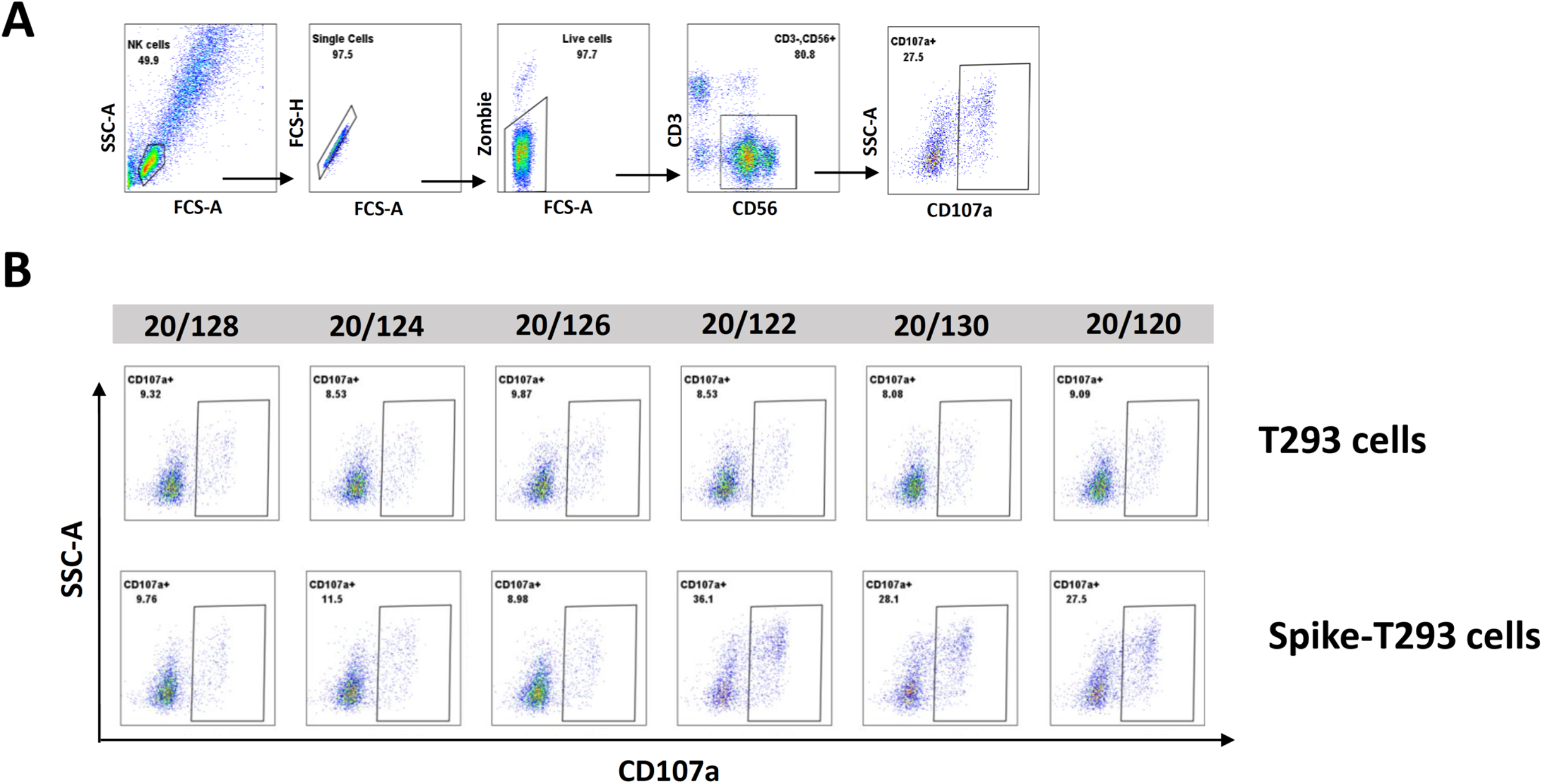
Antibody-dependent cellular cytotoxicity (ADCC) assay. (**A**). Representative gating strategy. NK cells were identified by lymphocyte morphology, singlet, live cell and CD3^-^ CD56^+^. The ADCC activity was defined based on NK cell degranulation (CD107^+^). (**B-C**) ADCC activity of NK cells induced by incubation with COVID-19 reference plasma panel (NIBSC 20/120, 20/124, 20/124, 20/126, 20/128, 20/130) obtained from the NIBSC in the presence of 293T-spike cells or 293T control cells. The experiment was performed in duplicate.

**Figure S7.**
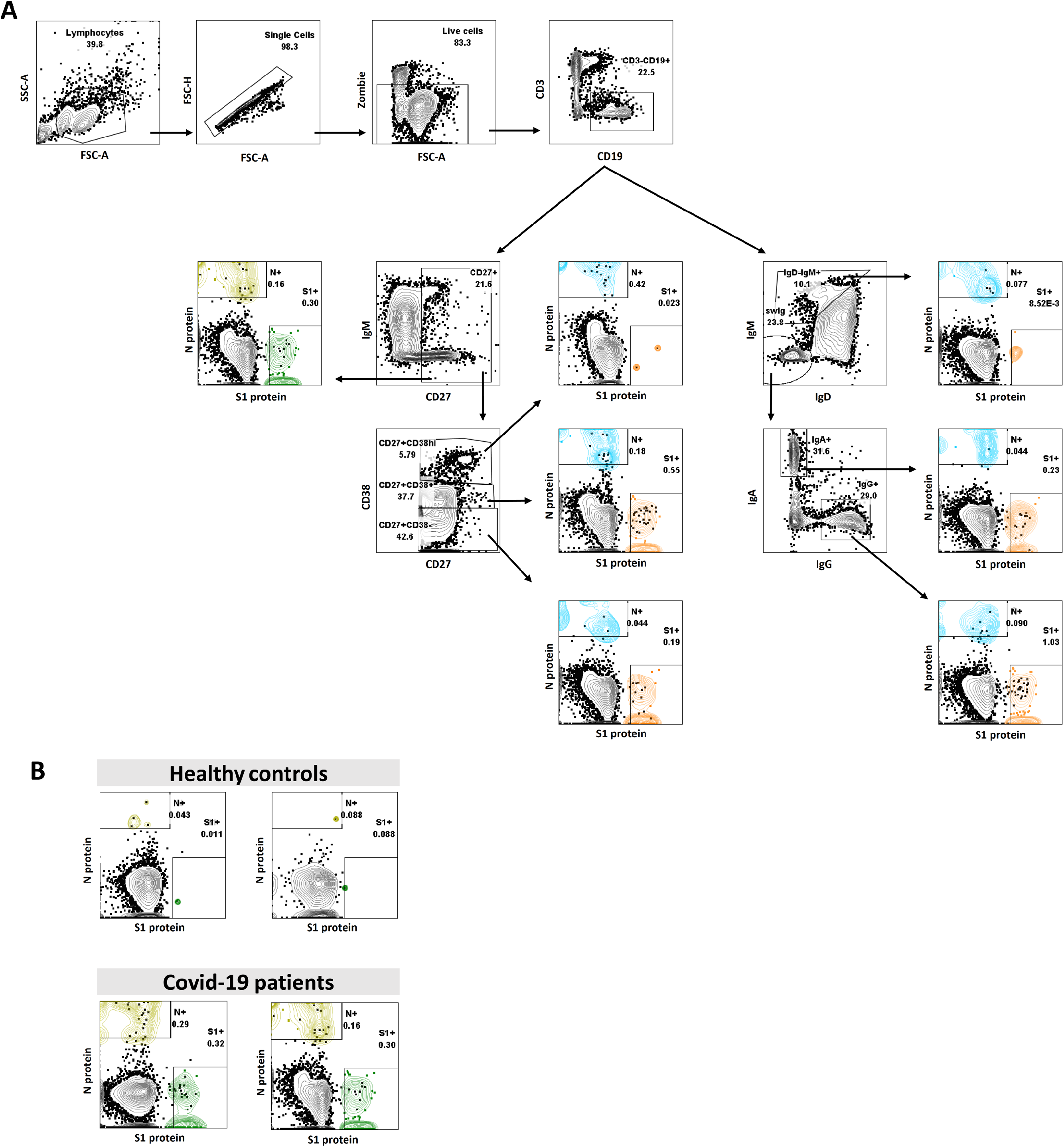
Representative gating strategy used to define antigen-specific B cells. (**A**) B cells were defined from the gates of morphology of lymphocytes, singlets, viability and CD3^-^CD19^+^. (**B**). Total B cells (CD19^+^) were further gated for B cell subsets including resting memory B cells (CD27^+^CD38^-^), activated memory B cells (CD27^+^CD38^+^) and plasma blasts (CD27^+^CD38^hi^). S1-specific B cells and N-specific B cells were determined. (**C**) Total B cells (CD19^+^) were further gated for B cell classes including non-class-switched mature cells (IgD^-^ IgM^+^) and class-switched IgM^-^IgD^-^IgA^+^ and IgM^-^IgD^-^IgG^+^ cells. The S1-specific B cells and N-specific B cells were determined. (**D**) Representative image comparing S1- and N-specific B cells in healthy controls and SARS-CoV-2-infected patients 6-9 months post infection.

**Figure S8.**
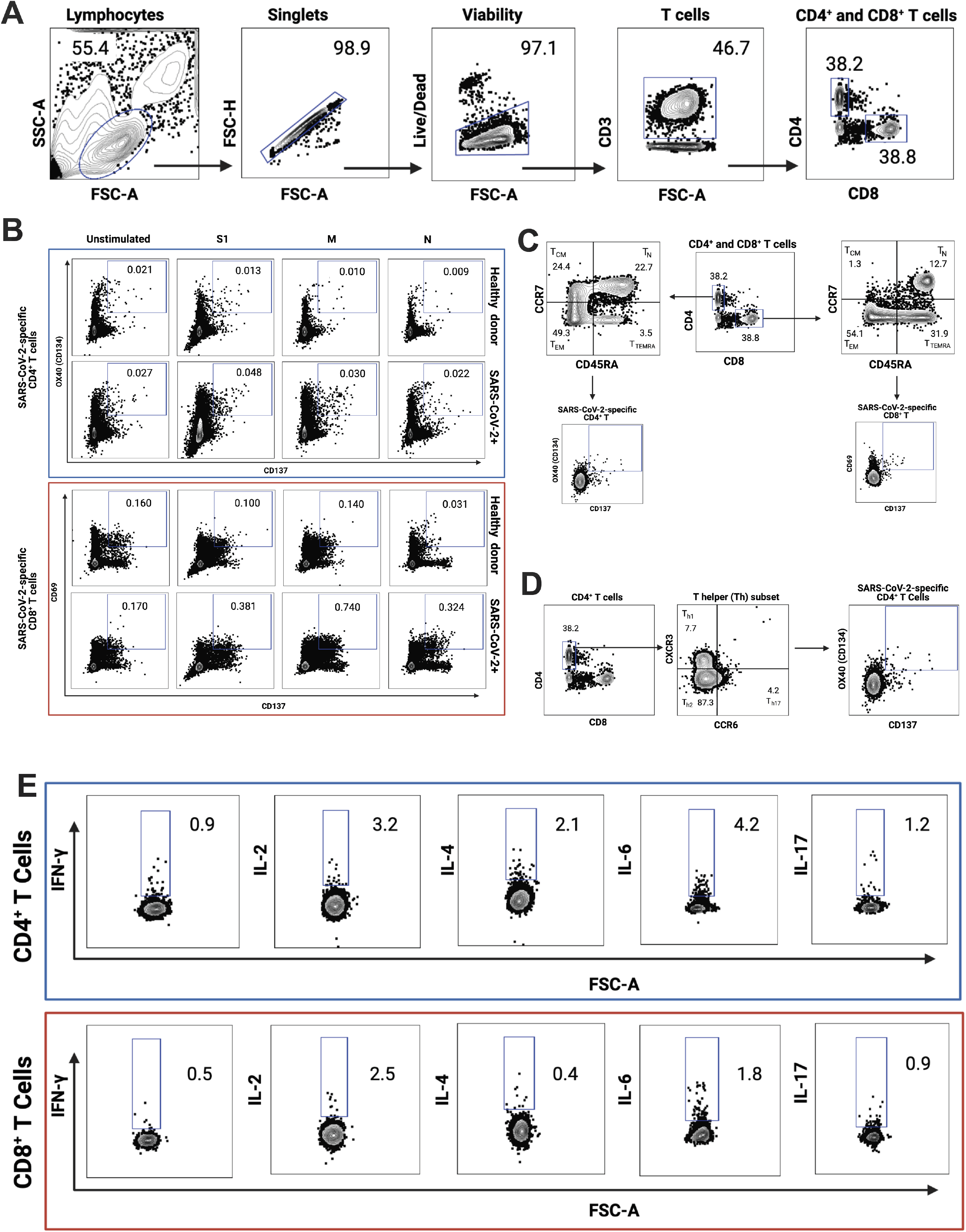
Representative gating strategy used for the T cells assays. **(A)** T cells were defined from the gates of morphology of lymphocytes, singlets, viability and CD3^+^. **(B)** Gating strategy used in the CD4^+^ and CD8^+^ T cell activation induced marker (AIM) assay to assess the SARS-CoV-2-specific CD4**^+^** and CD8^+^ T cells after overnight stimulation with S, M and N peptide pools. Representative image comparing S-, M- and N-specific CD4^+^ and CD8^+^ T cells in a healthy control and a SARS-CoV-2-infected patient 6-9 months post infection. **(C)** Gating strategy used in the CD4^+^ and CD8^+^ T cell activation induced marker (AIM) assay to assess the SARS-CoV-2-specific memory CD4**^+^** and CD8^+^ T cell subsets. Distribution of central memory (TCM), effector memory (TEM), and terminally differentiated effector memory cells (TEMRA) among total SARS-CoV-2–specific T cells. **(D)** Gating strategy used in the CD4^+^ T cell activation induced marker (AIM) assay to assess SARS-CoV-2–specific T helper (Th) subsets. **(E)** Gating strategy used in the CD4^+^ and CD8^+^ T cell intracellular staining assay to assess the cellular cytokine profile after 6 hours stimulation with S, M and N peptide pools.

**Figure S9.**
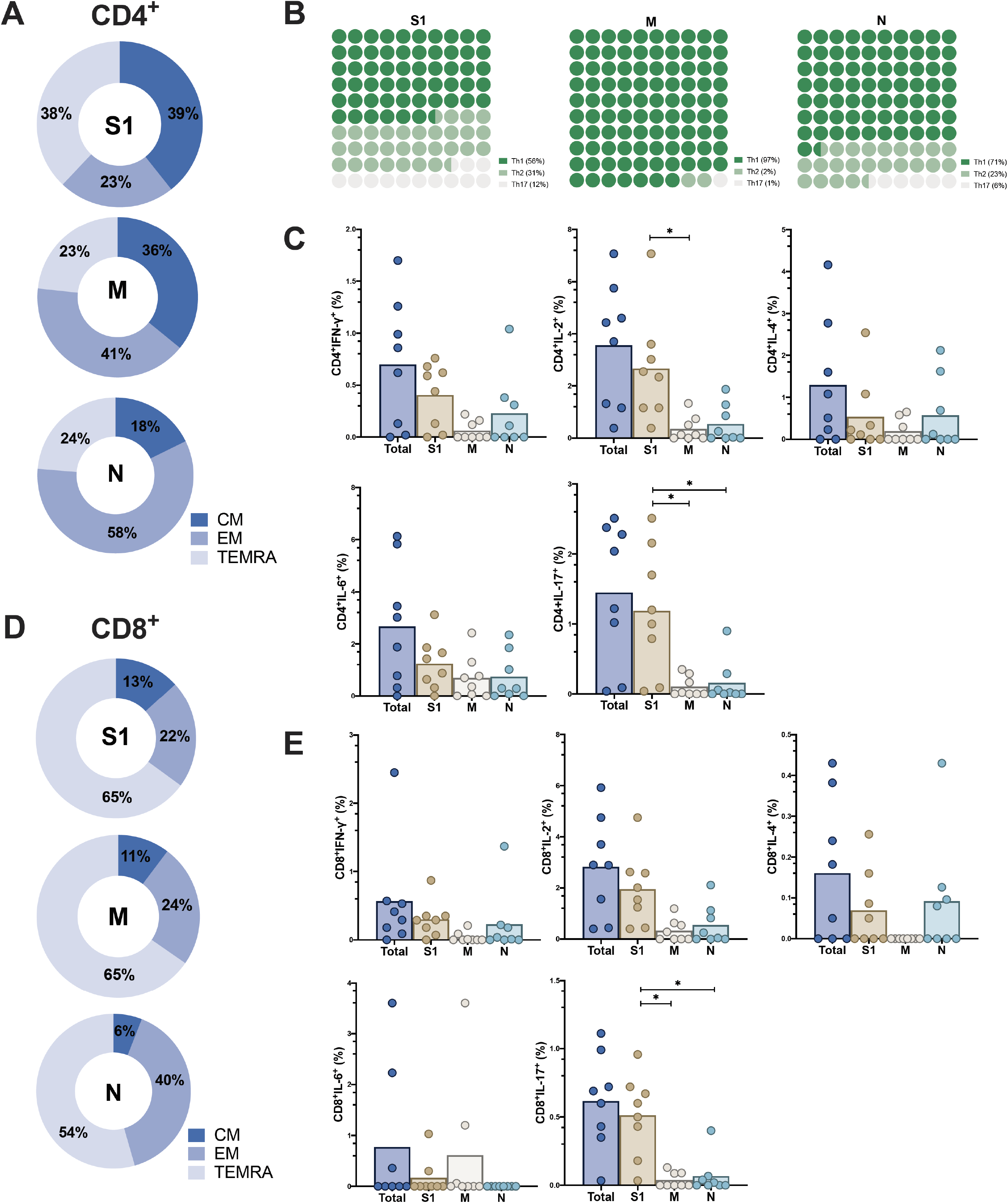
Characterization of SARS-CoV-2-specific T Cells. **(A)** Distribution of central memory (TCM), effector memory (TEM), and terminally differentiated effector memory cells (TEMRA) CD4^+^ T cells targeting different proteins of SARS-CoV-2 after overnight stimulation with different peptide pools. **(B)** The CD4^+^ Th differentiation, targeting different proteins of SARS-CoV-2, after overnight stimulation with different peptide pools. **(C)** Cytokine profile of CD4^+^ T cells after 6 hours stimulation with S, M and N peptide pools. **(D)** Distribution of central memory (TCM), effector memory (TEM), and terminally differentiated effector memory cells (TEMRA) CD8^+^ T cells targeting different proteins of SARS-CoV-2, after overnight stimulation with different peptide pools. **(E)** Cytokine profile of CD8^+^ T cells after 6 hours stimulation with S, M and N peptide pools.

**Figure S10.**
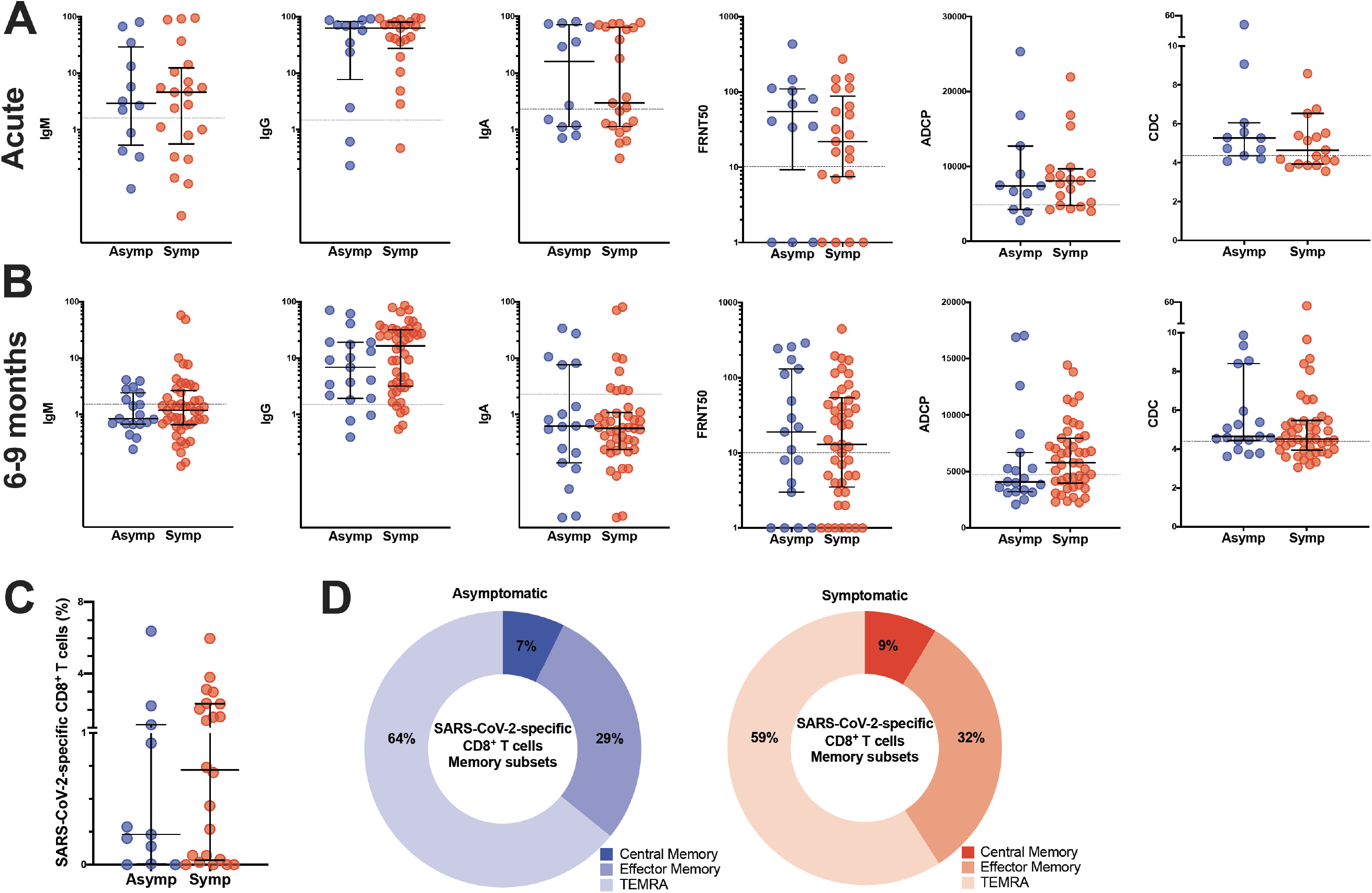
Comparison of immune parameters in asymptomatic and symptomatic individuals. Comparison of anti-S antibody titers, FRNT50 titers and anti-S mediated effector functions in the **(A)** acute phase and **(B)** late convalescent phase after infection. **(C)** Comparison of the frequency of total SARS-CoV-2-specific CD8^+^ T cells after overnight stimulation with peptide pools in asymptomatic individuals (asymp; n=11) and symptomatic patients (symp; n=22) at late convalescence. **(D)** Comparison of CD8^+^ T cell memory phenotype between asymptomatic individuals (asymp; n=11) and symptomatic patients (symp; n=22).

**Table S1:**
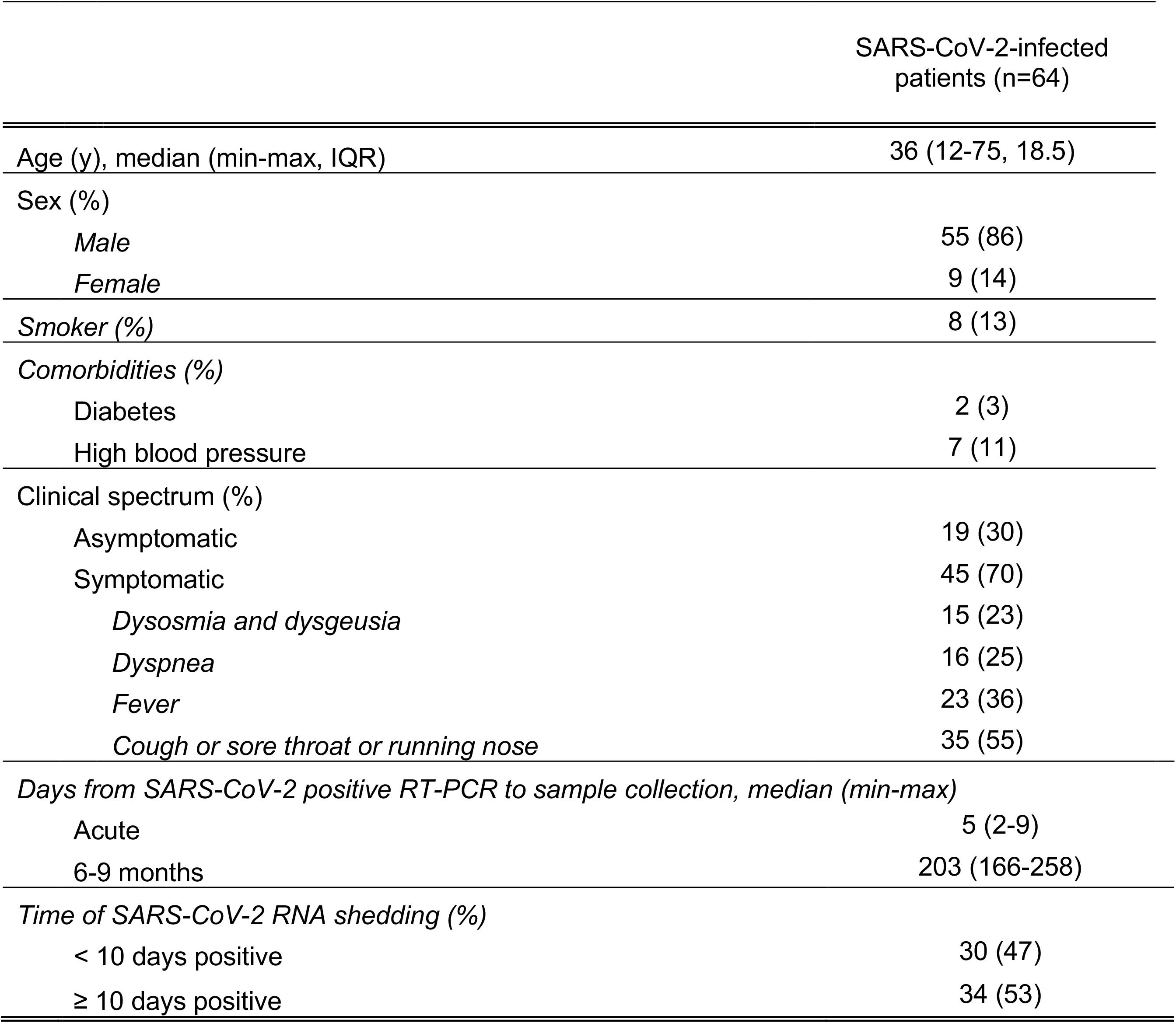
Cohort description.

**Table S2:**
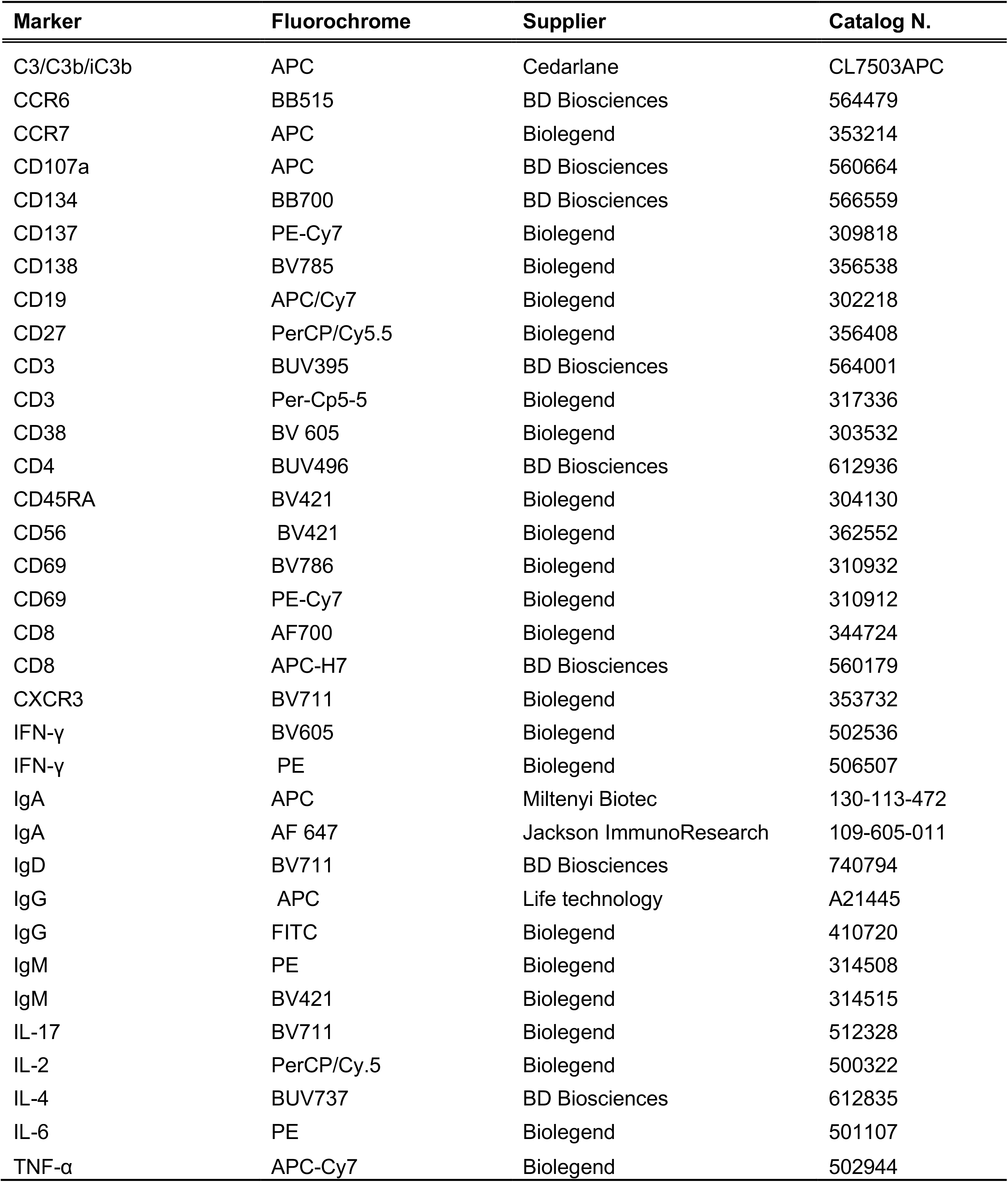
Monoclonal antibody list.

| Marker | Fluorochrome | Supplier | Catalog N. |
| --- | --- | --- | --- |
| C3/C3b/iC3b | APC | Cedarlane | CL7503APC |
| CCR6 | BB515 | BD Biosciences | 564479 |
| CCR7 | APC | Biolegend | 353214 |
| CD107a | APC | BD Biosciences | 560664 |
| CD134 | BB700 | BD Biosciences | 566559 |
| CD137 | PE-Cy7 | Biolegend | 309818 |
| CD138 | BV785 | Biolegend | 356538 |
| CD19 | APC/Cy7 | Biolegend | 302218 |
| CD27 | PerCP/Cy5.5 | Biolegend | 356408 |
| CD3 | BUV395 | BD Biosciences | 564001 |
| CD3 | Per-Cp5-5 | Biolegend | 317336 |
| CD38 | BV 605 | Biolegend | 303532 |
| CD4 | BUV496 | BD Biosciences | 612936 |
| CD45RA | BV421 | Biolegend | 304130 |
| CD56 | BV421 | Biolegend | 362552 |
| CD69 | BV786 | Biolegend | 310932 |
| CD69 | PE-Cy7 | Biolegend | 310912 |
| CD8 | AF700 | Biolegend | 344724 |
| CD8 | APC-H7 | BD Biosciences | 560179 |
| CXCR3 | BV711 | Biolegend | 353732 |
| IFN-γ | BV605 | Biolegend | 502536 |
| IFN-γ | PE | Biolegend | 506507 |
| IgA | APC | Miltenyi Biotec | 130-113-472 |
| IgA | AF 647 | Jackson ImmunoResearch | 109-605-011 |
| IgD | BV711 | BD Biosciences | 740794 |
| IgG | APC | Life technology | A21445 |
| IgG | FITC | Biolegend | 410720 |
| IgM | PE | Biolegend | 314508 |
| IgM | BV421 | Biolegend | 314515 |
| IL-17 | BV711 | Biolegend | 512328 |
| IL-2 | PerCP/Cy.5 | Biolegend | 500322 |
| IL-4 | BUV737 | BD Biosciences | 612835 |
| IL-6 | PE | Biolegend | 501107 |
| TNF-α | APC-Cy7 | Biolegend | 502944 |

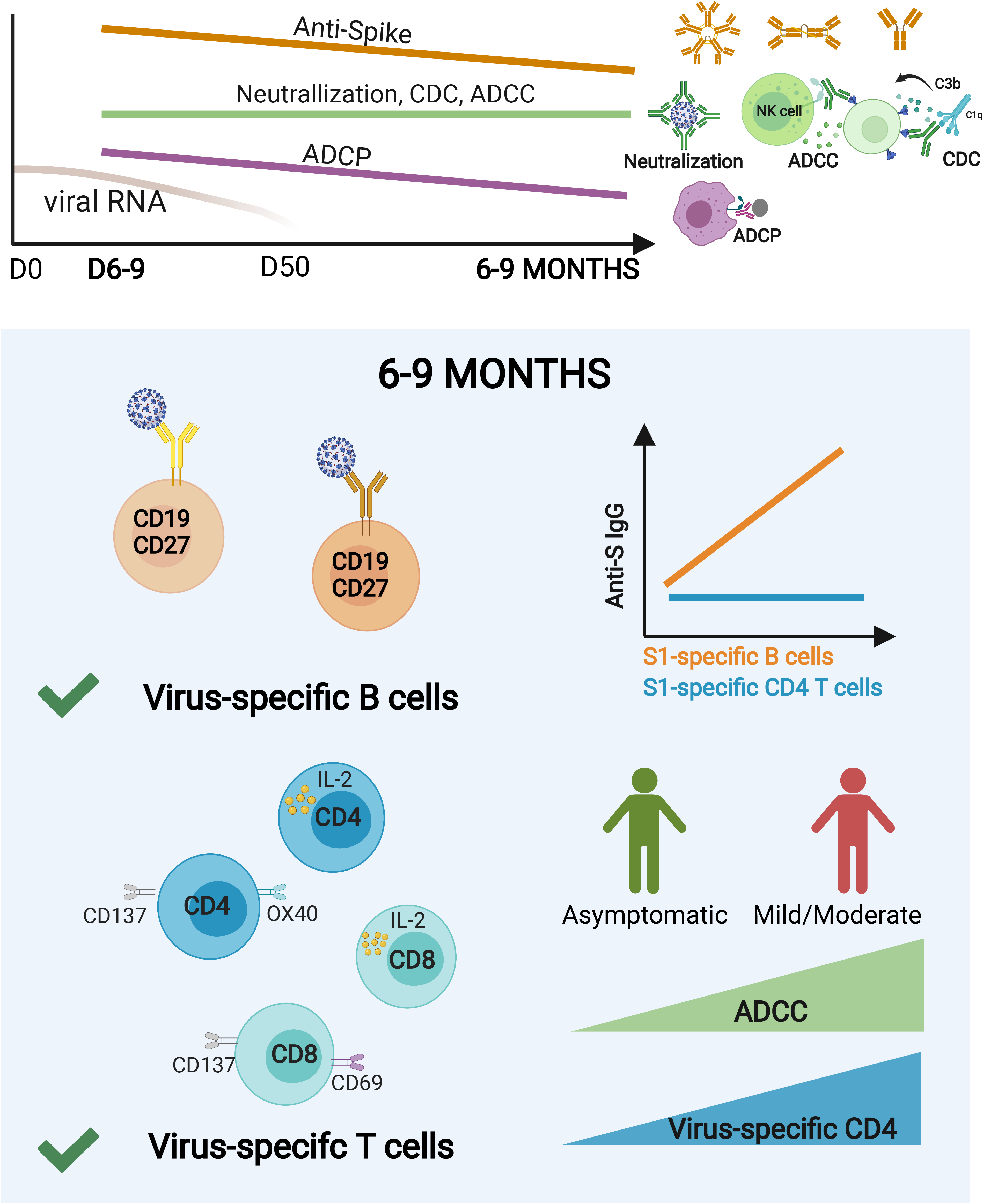

